# Structural basis of caspofungin inhibition of *Nakaseomyces glabratus* Fks1

**DOI:** 10.64898/2026.10.21.760291

**Authors:** Md Shamiul Islam Rasel, D Raghav, Fahrin Jahan Suchana, Rodolpho Souza Amado De Carvalho, TM Tomasiak

## Abstract

The Fks1 catalytic subunit of fungal glucan synthase crosslinks UDP-glucose into β −1,3-D-glucan in the fungal cell wall and is a target for the echinocandin class of antifungal drugs. Here, we present the resting state of Fks1 from *Nakaseomyces glabratus* (*Ng*Fks1) in both apo and caspofungin-bound states. Caspofungin binds the extracellular glucan exit portal of Fks1 adjacent to the product β -1,3-D-glucan in a position similar to that reported for *S. cerevisiae* Fks1. Unlike *S. cerevisiae* Fks1, no Rho1 binding is present, but the glucan channel opening is in an open-like state at the resistance hotspot regions. However, the lower gate is closed, suggesting glucan is trapped from a previous round of catalysis. Our results show that caspofungin can bind in a mode in the absence of Rho1 in *N. glabratus* and point to possible species-specific differences in substrate translocation, inhibition, and resistance mechanisms.

## Main Text

Fungal infections are an increasing public health threat worldwide, contributing to 3.8 million deaths per year ^1^. This affects immunocompromised patients, hospitalized populations, and individuals with limited access to rapid diagnosis and antifungal therapy ^2–4^. New resistant strains and innately resistant species, such as *Nakaseomyces glabratus*, have emerged in several continents in the last few decades ^5–8^. These pathogens have almost completely nullified the first line of common orally available antifungals, the azoles^9,10^. In the early 2000s, new classes of IV-administered antifungals derived from natural products called the echinocandins were released with the drug caspofungin (CFN) and followed by micafungin and anidulafungin ^11–19^. They showed high activity against many fungal species with low side effects ^20–23^ and have emerged as first-line therapy for many forms of invasive candidiasis ^18,24^.

Echinocandins target the catalytic subunit, Fks1, of the β -1,3-glucan synthase (GS) complex, the critical step in cell wall biosynthesis in several fungi ^25–27^. Fks1 was first identified in *Saccharomyces cerevisiae* and polymerizes UDP-glucose as a substrate, connecting glucose monomers to a growing glucose chain ^25,28^. Cryo-EM structures of Fks1 alone ^21,23^ and Fks1 in complex with the regulator subunit Rho1 revealed that Fks1 makes a large multidomain architecture that organizes around a glucan pathway connecting the cytoplasmic catalytic subunits to outside the membrane. This pathway resembles glucan synthases from other organisms ^29–31^. In several organisms, including *Candida albicans*, *N. glabratus*, and *Candidozyma auris*, resistance has been observed both *in-vitro* and in clinical settings ^32–36^. Mutants tend to cluster onto the membrane-facing surface of the transmembrane region and are classified into “hotspot” regions called hotspot-1 (625-633), hotspot-2 (1340-1344), and hotspot-3 (676-686), which are highly conserved regions ^32,34^.

The echinocandin mechanism and hotspot resistance mechanisms have been open questions since echinochandins were introduced ^37^. Echinocandins are large (> 1 kDa), amphipathic lipopeptides made of a 6-membered cyclic peptide headgroup and an N-linked alkyl tail^15^. Because of their amphiphilic nature, drug binding was presumed to be in the lipid bilayer, but various suggested models disagreed if they bind Fks1 directly or indirectly induce changes that alter activity ^38–41^. Early attempts to trap caspofungin failed ^21,42,43^. A recent cryo-EM study with the Fks1-Rho1 complex trapped Fks1 mid-catalysis show that the catalytic site is separated from the proposed glucan tunnel by ∼30 Å and showed the echinocandin inhibition mechanism proceeds through the formation of a glucan-drug ternary complex where the drug binds near the exit gate of the newly synthesized glucan polymer ^44^. Remarkably, caspofungin makes minimal contact with Fks1 itself and instead forms numerous contacts with the glucan chain itself. Importantly, this structure revealed that clinical hotspot mutants cluster near the caspofungin site and are components of the glucan exit gate.

In this work, we have determined the structure of pathogenic *N. glabratus* Fks1(NgFks1) in two new states – a post-catalytic apo state and a post-catalytic caspofungin-bound state. Both states capture glucan density in the exit gate; hence, we call it the post-catalytic state. The caspofungin-bound structure shows a striking resemblance in the hotspot region to *S. cerevisiae* Fks1 bound to the drug ^44^, despite lacking Rho1. Comparisons to the apo state suggest that NgFks1 may exhibit species-specific differences or has been purified in a new state that allows drug binding.

## Results

### Structure determination of NgFks1

We determined the structures of Fks1 from the pathogen *N. glabratus* in the apo state and a caspofungin-bound state with the central glucan pathway blocked by glucan in both. We extracted Fks1 from yeast membrane using lauryl maltose neopentyl glycol (LMNG) and cholesteryl hemisuccinate (CHS) at a deliberately low concentration to maximize the retention of endogenous lipids (0.5% LMNG compared to typical 1-1.5%). The protein was then purified using nickel affinity chromatography, where the buffer was exchanged with 0.06% glyco-diosgenin (GDN) followed by size-exclusion chromatography (SEC) with the buffer containing 0.02% glyco-diosgenin (GDN).

We performed single-particle cryo-EM reconstruction, which generated 3D maps of the NgFks1 apo state at 3.47 Å (**Fig. 1*B***) and the NgFks1 caspofungin-bound state at 3.32 Å (**Fig. 1*C***). Model building revealed that Fks1 contains 15 well-resolved transmembrane helices **(Fig. S3*A-B*)**, which we numbered according to the new numbering scheme proposed for *S. cerevisiae* Fks1 (ScFks1) ^44^. The extracellular density and GT and AC domains were both well resolved, except for substrate-binding loops in GT, which were also disordered in the *S. cerevisiae* cryo-EM structures but have been resolved in newer structures purified in CHAPS detergent. TM1-6 makes the first cluster that flanks the catalytic core GT domain, while the second, larger cluster includes TM 7-15 and harbors the major architecture of the glucan translocation channel (**Fig. 1*A***). Two conserved disulfide bonds have also been resolved in the extracellular loop - C644-C655 (TM5 and TM6 connecting loop) and C1315-1332 (TM7 and TM8 connecting loop).

**Fig. 1.**
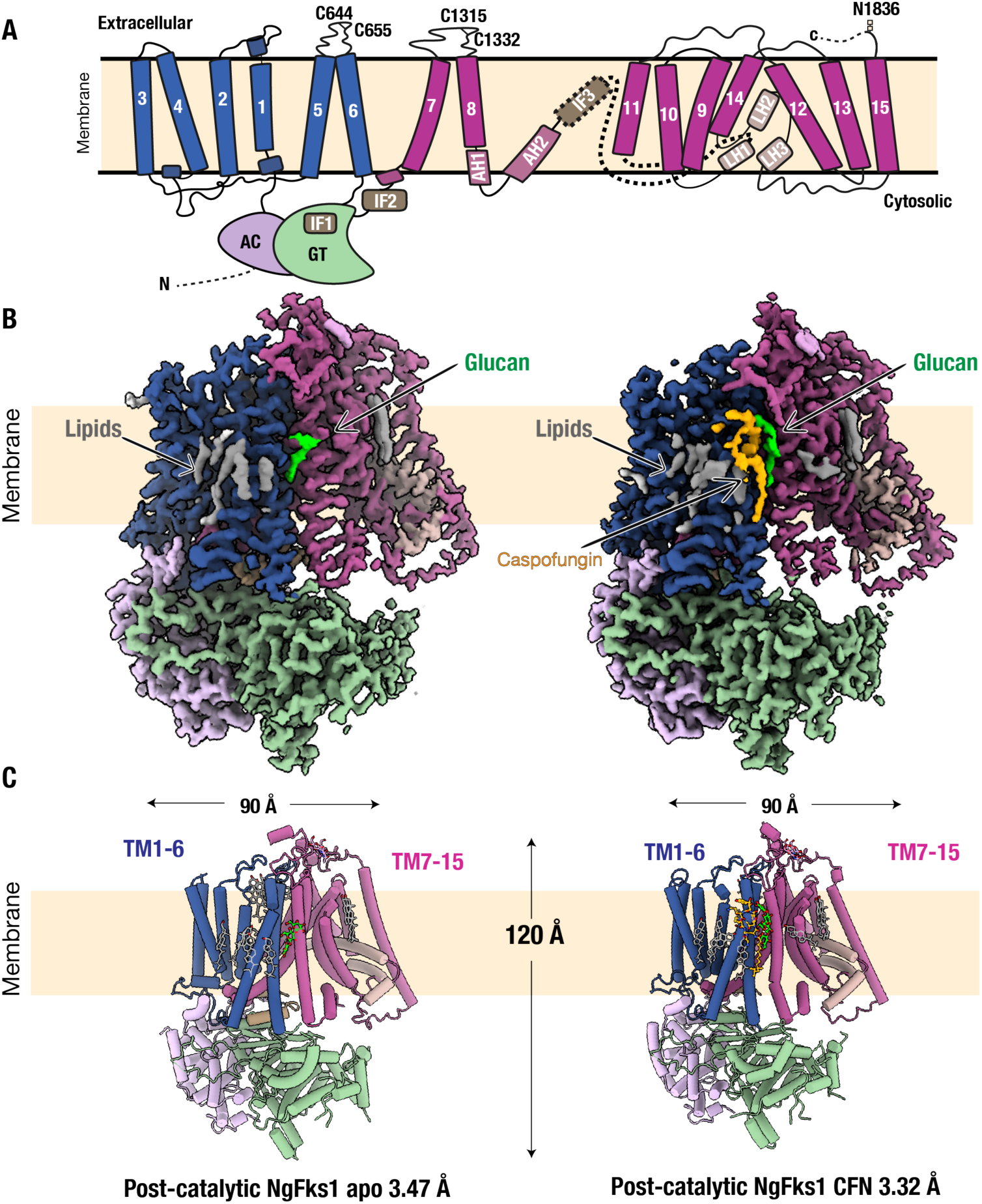
Architecture and structure of *N.glabratus* Fks1. **(A)** Topology diagram showing the layout of domains in NgFks1:AC domain (lavender), TM helices 1-6 (blue), TM helices 7-15 (purple) and GT domain (light green). Conserved elements shown include IF1, IF2, IF3 (tan), LH1, LH2, LH3 (light tan) and AH1 and AH2 (light purple). The dotted line indicates disordered IF3. Two conserved disulfide bonds are shown between C644-C655 and C1315-C1332. Conserved N-linked glycosylation is shown at N1836. **(B)** Cryo-EM map of post-catalytic NgFks1 apo state at 3.47 Å and NgFks1 caspofungin-bound state at 3.32 Å. Caspofungin is shown in gold, lipids in grey, glucan densities in lime green, the membrane is represented as a beige slab. **(C)** Cryo-EM model of NgFks1 apo and caspofungin-bound state.

### Caspofungin and glucan-bound state

The caspofungin-bound state resulted from caspofungin incubation of the purified protein sample. We incubated the purified Fks1 with 200 µM caspofungin at 20°C for 1 hour before sample freezing. Examination of the density revealed a structure in the TM region similar to the Rho1-CFN-bound ScFks1 structure (pdb: 9pe1), except for a closed TM8 glucan gate spanning residues 1356-1376 ^44^. However, unlike previous structures of Fks1 alone, strong density in the glucan pocket region remains visible, enabling the modeling of 4 subunits of the glucan polymer.

Strong density of caspofungin was immediately evident in the cryo-EM maps and was missing in our apo structure (**Fig. 2*A***), consistent with the position observed in the Rho1-CFN-bound ScFks1 structure ^44^. The lipidic tail P1 (Fig. S2) binds in the same groove along the transmembrane region but is slightly shifted down towards the lipid bilayer (**Fig. 2*B***). The entire peptidic head could be modelled in a conformation mostly consistent with the Rho1-CFN-bound ScFks1 structure but with slight differences in segments that do not bind glucan (**Fig. 2*C***). While the peptidic headgroup Thr (P2) and 3OH-Pro (P3) positions are similar, the (OH)2-hTyr (P4), OH-Orn (P5), 4OH-Pro (P6) and ethylenediamine (P7) groups lie flat against Fks1 as opposed to their large surface of engagement with glucan in the Rho1-CFN-bound ScFks1. However, this likely reflects the smaller amount of glucan density (4 subunits modelled versus 17 in ScFks1) and the TM8 opening in ScFks1. In our structure, the different position of TM8 shifts the position of glucan and seemingly caspofungin along with it, where otherwise there would be a clash with our observed glucan.

**Fig. 2.**
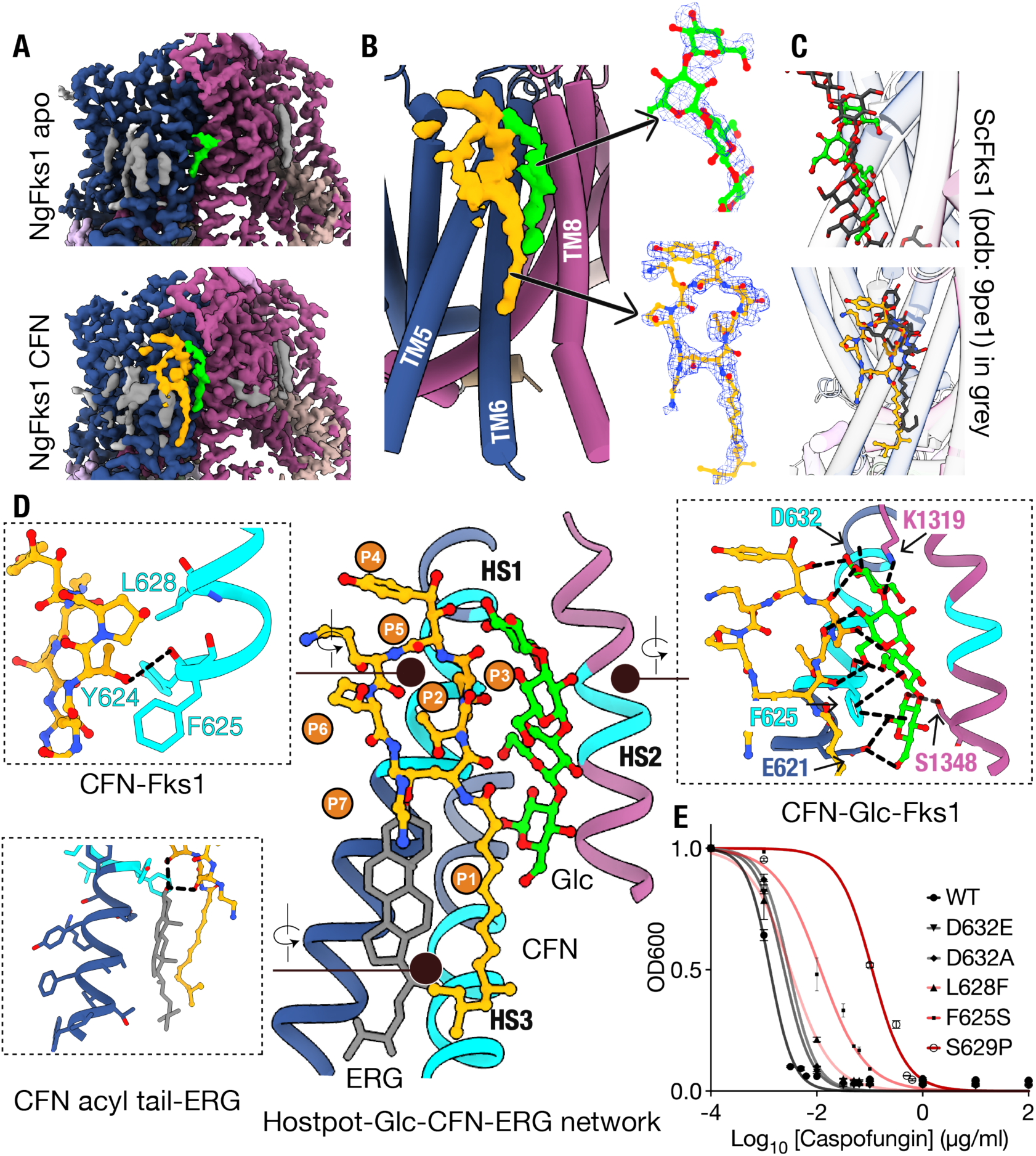
Caspofungin binding with NgFks1. **(A**) Cryo-EM density comparison between NgFks1 Apo and CFN-bound state, showing additional drug density in gold. **(B)** A close-up view of the drug-binding region shows that caspofungin binds on the interface of transmembrane helices 5, 6, and 8 near the translocating glucan fragments. **(C)** Comparison of the caspofungin (yellow) and glucan fragment (green) positions to the published Rho1-CFN-bound ScFks1 complex (shown in dark grey - pdb: 9pe1). **(D)** A close-up view of the interaction network between caspofungin, glucan fragment, hotspot adjacent residues and nearby sterol. H-bonds are shown with the dotted line. The caspofungin interacts with hotspot 1 by H-bond and hydrophobic interaction (top left). and the glucan interacts with CFN through multiple H-bond interactions, with Fks1 via H-bond with E621, S1348, D632 and K1319 and hydrophobic interaction with F625 (top right). The bound sterol shows H-bond and hydrophobic contacts with the CFN (bottom left). **(E)** Survival assays of hotspot mutants. The growth was measured by OD600 and normalized to the no-drug control for each strain. Each data point is a representation of 3 biological replicates.

Instead, caspofungin makes extensive contacts with an adjacent bound sterol (**Fig. 2*D*, bottom-left inset)**. The caspofungin lipid tail directly packs against the ergosterol through hydrophobic and H-bond contact. This implies that caspofungin can readily be bound to the less extensive glucan assemblies in Fks1 and only engages fully with the active state and a fully extended glucan chain.

Similar to caspofungin binding, glucan binding also recreates many of the contacts seen in the Rho1-CFN-bound ScFks1 structure but is more discontinuous. Glucan densities appear to occlude the exit gate at multiple levels – lower gate around residues E621, D677 and I1351, mid gate residues around F625, S1348 and L1347 and upper gate residues S629, R1340 and R1343. The glucan fragments are stabilized by several key interactions, with residues E621, F625, S1348, D632 and K1319 (**Fig. 2*D*, right inset).** Extensive H-bond interactions are observed between the backbone carbonyls along P1-P4 groups of CFN and the glucan polymer (**Fig. 2*D*, right inset)**, which likely contribute to the major stabilizing forces that bind the CFN in the glucan exit gate. At the base of the glucan chain, the glucan pathway is closed off by the positioning of W681, a highly conserved residue in hotspot 3 that has been proposed to form the lower gate of the glucan path. Notably, similar glucan-like density is also present in the reported enfumafungin-bound ScFks1 structure, but left unmodeled ^43^.

In addition to interactions made between portions of the glucan exit gate, bound sterols, and the glucan polymer, caspofungin makes only a few direct contacts with Fks1(**Fig. 2*D***). The most pronounced are in the 10,12-dimethylmyristoyl lipid tail, which binds a groove directly adjacent to the glucan tunnel with a large interaction to the hydrophobic surface of TM5 and TM6, made of residues F625, W681, V684, and V688 bordered by interactions with ergosterol. The P2 and P3 make hydrophobic contacts with Y624, F625, and L628 and P2 crucially makes H-bonds to the main-chain carbonyls at residue Y624 of helix TM5 (**Fig. 2*D*, top-left inset).**

The glucan fragments make contacts along the P1-P4 surface of caspofungin. Interestingly, the majority of these contacts are made between the main chain carbonyl of caspofungin and the hydroxyl groups of the glucan chain.

### Caspofungin binding

We performed a Microscale Thermophoresis (MST) binding assay to confirm caspofungin binding in GDN-purified NgFks1 without addition of rho1. The assay shows concentration-dependent binding of caspofungin with a Kd of ∼360 µM (Fig. S4). The value coincides with the binding assay of NgFks1 with enfumafungin, a triterpenoid inhibitor ^45^, but is likely higher than in native lipids because of the large GDN micelle in both cases.

For functional characterization of the clinical hotspot 1 and echinocandin binding sites, we performed cell survival assays. We generated an Fks1 knockout line in the BJ2168 strain of *S. cerevisiae*, where Fks2 was inhibited using Fk506 because of some functional redundancy ^21^. Then we complemented the cells with NgFks1 transformation, which showed robust growth, meaning NgFks1 can substitute for the ScFks1 activity. We created 5 hotspot clinical mutants ^44^ (S629P, F625S, L628F, D632A, D632E) on hotspot 1 and performed a growth assay (**Fig. 2*E***). The hotspot mutants showed high caspofungin resistance that is consistent with clinical MIC50 data ^34^. The most pronounced resistance came from S629P, which increased nearly 100-fold compared to WT. The F625S confers the second strongest resistance, L628F shows intermediate resistance, and D632E/A confers the mildest resistance to caspofungin.

### Exit gate remodeling

Our NgFks1 apo structure lacked density for caspofungin, as expected, but surprisingly showed limited density for glucan in the same position as our caspofungin and the Rho1-CFN-bound ScFks1 complex (Fig. S5*A,B*). The interactions with E621, D677, F625, and S1348 stabilize the bound glucan (Fig. S5*B*).

The trapped glucan fragment and caspofungin induce a cascade of side-chain, backbone and helical changes across both the TM5 and TM8 helices, where hotspots 1, 2 and 3 are located. We observed a helix break at TM8 region in our drug-bound structure when compared to the apo. Notably, the break happens at a similar location where the glucan passes through the lower gate of the exit tunnel (**Fig. 3*A***). This suggests CFN binding itself drives Fks1 to an active-like state, or the initial transition to the active-like state might influence caspofungin binding.

**Fig. 3.**
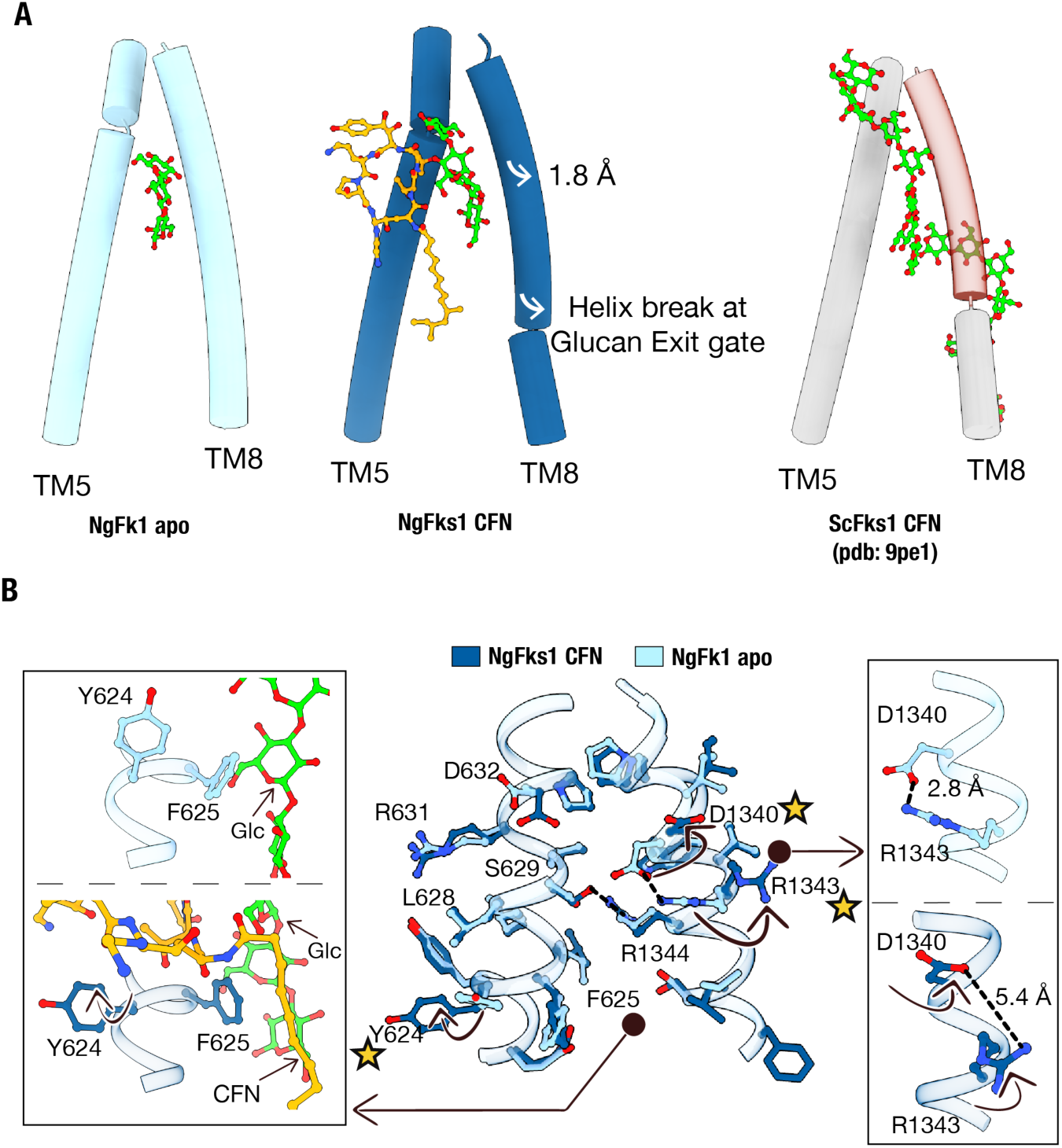
Comparison of the glucan exit pathway near hotspot 1 and 2 regions. **(A)** Exit gate comparison across NgFks1 apo, NgFks1 CFN and ScFks1 CFN (pdb: 9pe1) showing a helix break at the lower TM8 exit gate and a 1.8 Å shift at the upper TM8 gate in the caspofungin-bound state. TM8 moves out at a similar location to the helix break in the ScFks1 CFN. **(B)** Side chain comparison between NgFks1 apo (sky blue) and NgFks1 CFN-bound state (dark blue) shows the altered conformation of Y624, R1343, and D1340. The changes are shown with stars. Close-up views of the altered residues are shown in insets.

Compared with each other, the apo and CFN-bound structures show side-chain rearrangements in residues Y624, D1340, and R1343 (**Fig. 3*B***) that coincide with a similar but smaller glucan density in the apo structure than in the CFN-bound structure. In the CFN-bound structure, Y624 undergoes a 90-degree outward rotation to provide space for the placement of the drug’s P2 group (**Fig. 3*B*, *left inset*)**. Our apo structure captures a salt bridge not observed in other reported Fks1 structures, formed between D1340 and R1343 within the TM5 and TM8 cavity. In our CFN-bound structure, the TM5 and TM8 cavity swings outward, breaking this salt bridge (**Fig. 3*B*, right inset)**. We propose that this transition may allow the R1344 movement, accompanied by TM8 movement, observed in the long-glucan CFN-bound ScFks1 (pdb: 9pe1).

To understand the gating dynamics, we compared the key gating and resistance hotspot residues Y624, F625, R1344, R1343 and D1340 across different states of Fks1 from our structures to recently published *S. cerevisiae* structures (**Fig. 4A**). We defined side chain rotamer positions as “in” or “out” based on their position relative to the glucan channel between TM5 and TM8.

**Fig. 4.**
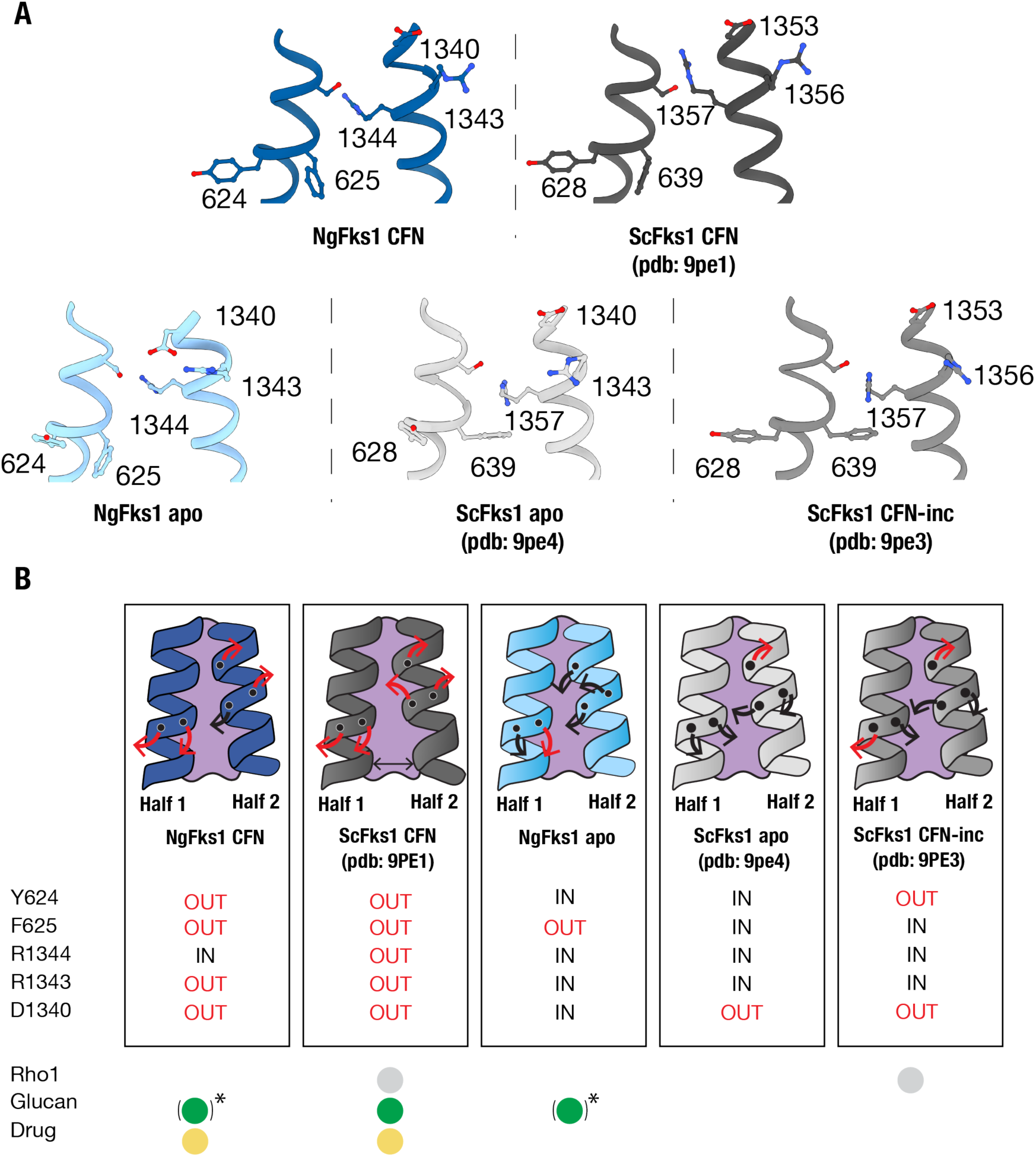
Summary of hotspot 1 and hotspot 2 side chain changes in different states of Fks1. **(A)** Side-by-side hotspot site key residue (Y624, F625, R1344, R1343 and D1340) comparison of NgFks1 CFN-bound post-catalytic state (dark blue) and ScFks1 CFN-bound active state (dark grey) (pdb: 9pe1) on the top panel. Comparison of NgFks1 apo post-catalytic state (sky blue), ScFks1 apo (light grey) (pdb: 9pe4) and ScFks1 CFN-treated state in tin grey (pdb: 9pe3) in the bottom panel. **(B)** Black arrows indicate inward movement of the residues toward the glucan channel; red arrows indicate outward movement. S629 is not shown and is identical across all structures. Pdb: 9pe1 is the only one depicting an open glucan channel and R1344 rotamer movement is up. The * near glucan represents a minimal glucan fragment.

Interestingly, our NgFks1 CFN state shows an almost identical arrangement to the ScFks1-CFN state. The most notable exception is an upward shift of R1344 in ScFks1, which is absent in NgFks1 and may be tied to Rho1 binding and TM5 movement. This movement nevertheless maintains similar binding to partners, including S629. It differs from several apo states of ScFks1. The ScFks1 apo state (pdb: 9pe4) represents a basal state where all the gating residues are in closed conformation except D1340. The ScFks1 drug-incubated state (pdb: 9pe3) represents a CFN-treated ground state that nevertheless lacks CFN and glucan density that closely resembles the apo state, except for a Y624 outward movement.

Interestingly, the NgFks1 apo state didn’t show similarity with any of the states. Specifically, the outward-open F625 conformation resembles the open glucan conformation seen in 9pe1 structure and our NgFks1 CFN state, implying either that the gate is constitutively open in the pathogenic NgFks1 or by stabilizing a glucan fragment, we trapped F625 in an open conformation. It also reinforces the critical role of F625 as a gating residue or sensor for bound glucan, which was also reported in the recent study ^44^. The ScFks1 CFN state (pdb: 9pe1) shows the largest divergence from our apo structure, where all gating residues acquired the outward confirmation (**Fig. 4B**).

### The translocation channel

The inner glucan channel connects the catalytic site in the cytoplasm to the exit gate. The translocation channel residues remain largely unchanged between the two states (Fig. S6*A*). The only discernible difference is at K1069 (ScK1082) at the catalytic site, which directly anchors the non-reducing end of the growing glucan chain (pdb: 9pe1). The conserved DAN motif and the NED motif residues E1208 (ScE1221) and D1209 (ScD1222), which mediate UDP interactions, are also resolved and show only minor rotamer shifts between our two states. We performed liquid growth assays using strains expressing E837A, K1069A, D1209A, and K1248A NgFks1 (Fig. S6*B*). All mutants failed to grow comparably to wild type, reaching OD600 values similar to the empty-vector control. Together, the functional, structural, and sequence alignment (Fig. S1) data support NgFks1 as a mechanistic ortholog of ScFks1 in UDP-glucose catalysis and glucan chain propagation.

The channel also contains a core architecture formed by the catalytic loops IF1, IF2, and IF3, which have been resolved to varying degrees in previous structures and completely resolved in the Rho1-CFN-bound state of ScFks1 (9pe1). Similar to other Fks1-alone structures, most of the NgFks1 density we present, the IF loop density remains unresolved. The exceptions are NgFks1 apo, where IF1 is fully resolved and IF2 is partially resolved. Neither structure resolves IF3, which forms a critical wall for glucan (Fig. S6*C*).

### Ordered lipid pockets near the hotspot and catalytic sites

Several sterol groups form a cluster near the caspofungin binding site (Fig. S7*A*). One of these lipids (we call the scaffold lipid) makes one of the few contacts that hold caspofungin in its position. These lipids are stabilized by six aromatic residues (Y452, Y455, W479, F556, Y623 and Y624) along TM1, TM4, and TM5 (Fig. S7*B*). Together, they create a hydrophobic cage with a pi-electron-rich ceiling that sits beneath the important gating loop TM7-8.

We mutated these residues and performed a cell survival assay in the presence of CFN (Fig. S7*C).* Y623A and Y624A produced the strongest reduction in caspofungin sensitivity compared to WT; W479A was intermediate and Y455A, F556A, and Y452A produced progressively milder reductions. Overall, the growth profile was not as strong as seen in clinically resistant mutants (**Fig. 2*E***).

## Discussion

Our NgFks1 structure with caspofungin and glucan details a new, inactive Fks1 conformation. Glucan binding to hotspot regions shows a pattern similar to the ScFks1-CFN-Rho1 complex (pdb: 9pe1) ^44^ but with only a small glucan fragment and closed TM8 gate, likely because Rho1 is absent. This suggests an uncoupling of the lower and upper halves of the glucan channel. In addition, R1344 is readjusted down in the NgFks1 structure. We propose this state is CFN bound to an inactive or catalysis-terminated form of Fks1. For these reasons, we suggest that both our states are post-catalytic states that have been captured following extraction from cells and dissociation of Rho1.

Several factors suggest a drug-bound post-catalytic state. First, comparison of all Fks1 structures shows the closest resemblance is between our CFN-bound structure and the active Fks1-CFN-Rho1 structure from Baker’s yeast, which are almost identical in the hotspot regions with the exception of lower TM8 (**Fig. 4**). Second, in contrast, our apo structure resembles none of the previously determined structures either from our work or others. Glucan density is present, but the resulting pattern of hotspot residues rotamers differs from the CFN-bound structure, and all apo structures of Fks1. Finally, we and others show that NgFks1 can substitute for yeast Fks1 when overexpressed, suggesting it interacts with yeast Rho1 to become active. This cross-species activation by yeast presumably allowed the capturing of this state. During the preparation of this manuscript, a structure of NgFks1 was reported with the triterpenoid enfumafungin bound, which is in high agreement with our presented structure as well ^45^.

The caspofungin binding site here has implications for the drug activity and resistance. P1-P3 of CFN bind in a closely similar pose to that of ScFks1, with extensive and identical contacts to glucan, Fks1, and the scaffold lipids, including two sterols conserved in all Fks1 structures^21–23^. Notably, the CFN threonine and the 3-hydroxyproline (P2 and P3) both make contacts near the critical and most resistant S629 site, with an unwound helix in this position (**Fig. 2*D***). The rest of CFN P5-P7, however, is not in the same position and we infer that CFN binding to this state takes place in a slightly altered and less engaged drug conformation that nevertheless engages the same critical portions of the glucan gate (Fig. S8). CFN then likely requires a longer glucan chain to engage more fully.

In both the apo and drug-bound states, the glucan cavity is opened by movement of F625, which makes a binding surface for the lipid tail of CFN (**Fig. 2D**). Our apo structure also has this F625 position in the open state because of bound glucan. The state captured here contrasts with ScFks1, which could only be captured after binding Rho1. This could suggest possible differences between pathogenic NgFks1 to ScFks1, with a state susceptible to CFN even in conditions where Rho1 is disengaged. Nevertheless, this suggests states other than the fully activated enzyme can be targets for the drug and present potential new binding surfaces to target.

The differences between apo versus drug bound also help explain resistance mutant severity. The most severe, S629P, makes a carbonyl contact to CFN but also must flex in response to distortions from TM8 opening. CFN binds to TM6 around S629 to influence the helical break and also binds to Y624, F625 and L628, all of which are resistance hotspots. Thus, proline rigidifies this important surface and interferes with CFN-induced changes, binding, or both. Similarly, F625 must rotate 90° to accommodate the glucan molecule (9pe1) and ordered lipid^44^. The F625S mutant disrupts these key interactions.. Finally, D632 (ScD646) binds glucan in our structure and shows minimal resistance. Altogether, resistance is likely driven by secondary allosteric changes through this network in states we have not yet observed.

It is unclear why our structure binds CFN without Rho1 but could not in the *S. cerevisiae* ^44^. Reported Fks1 activity is highest in CHAPS detergent, which is required for *in vitro* Rho1 assembly 44. Our purification strategy instead involved using low amounts (0.5%) of extracting LMNG to preserve bound lipids. The closed glucan channel in NgFks1 suggests it was captured in an inactive state following a period of Fks1 assembly with endogenous Rho1 and activity during over-expression. Consistent with this model, our inner translocation regions show disordered IF loops in the catalytic site, consistent with other Fks1-only structures (pdb: 9pe4, 9pe5) ^44^. It is unclear if CHAPS or Rho1 is responsible for this difference.

Along with protein and glucan, a final binding interaction to CFN comes from two sterols, which are present in the *S. cerevisiae* structure as well ^22,23,44^. In NgFKS1, these sterols are themselves in contact with a large cluster of sterols in an aromatic pocket near caspofungin. Mutations in residues that destabilize these sterols lead to a resistance phenotype, presumably because they interfere with the caspofungin mechanism but still allow plasticity in the binding site. The sterol makes close contact with F625, which is a hotspot mutant, and notably binds enfumafungin in the *S. cerevisiae* Fks1 and CFN in NgFks1^43^. The lipid environment surrounding the glucan channel may play an important role in determining drug binding and resistance mechanisms.

## Materials and Methods

### Cloning, expression & purification of Fks1

The *N. glabratus* Fks1 gene was cloned into the pESC-Ura yeast expression vector after codon optimization, with N-terminal FLAG (DYKDDDDK) and C-terminal deca-histidine (10X His) tags (GenScript). The protein was expressed in S. cerevisiae strain BJ-2168 (Genotype: MATa prc1-407 prb1-1122 pep4-3 leu2 trp1 ura3-52 gal2). The Fks1-transformed yeast strain was cultured in uracil dropout YNB medium (0.67% wt/vol yeast nitrogen base without amino acids, 2% wt/vol glucose, and 0.08% wt/vol amino acid dropout mix without uracil) for at least 24 h at 30 °C. A secondary 750 ml culture was then inoculated with 15 ml of the primary culture and grown for an additional 24 h. The induction was done by adding 250 ml of YPG medium (1% wt/vol yeast extract, 1.5% wt/vol peptone, and 2% wt/vol galactose final concentration) and growing for 16-18 h at 30°C. The culture was then harvested by centrifugation at 4500 x g for 30 min at 4°C.

For purification, we followed a similar strategy as in our previous study ^46^. The harvested cells were lysed by bead beating using 0.5 mm glass beads in the lysis buffer containing 50 mM Tris-Cl pH 7.4, 300 mM NaCl and a protease inhibitor cocktail made of 1 µM Aprotinin, 2 µM Pepstatin, 10 µM Leupeptin and 0.5 mg/ml AEBSF. The lysate was subjected to centrifugation at 4000 rpm for 10 mins and the supernatant was collected for ultracentrifugation at 100,000 x g for 1.5 h at 4°C. The collected membrane pellet was solubilized in the resuspension buffer (50 mM Tris–Cl pH 7.0, 300 mM NaCl, 0.5% 2,2-didecylpropane-1,3-bis-β-D-maltopyranoside [LMNG]/0.05% cholesteryl hemisuccinate [CHS] supplemented with protease inhibitor as described above). After 4 hours of solubilization, the insoluble materials were separated by centrifugation at 35,000 x g for 30 mins at 4°C. The clarified supernatant was then applied to a 5 ml Ni-NTA immobilized metal affinity chromatography column (Bio-Rad), which was pre-equilibrated in Buffer A (50 mM Tris–Cl, 300 mM NaCl, 0.01% LMNG/0.001% CHS, pH 7.4). After a subsequent wash with a gradient of Buffer B (50 mM Tris–Cl, 300 mM NaCl, 500 mM imidazole, 0.01% LMNG/0.001% CHS, pH 7.4), the protein was eluted with 4 column volumes of 60% buffer B. The protein was concentrated and finally purified by size exclusion chromatography using a Superose 6 Increase 10/300 GL column (GE Healthcare) which was also pre-equilibrated with size exclusion buffer (50 mM Tris, 300 mM NaCl, pH 7.4) supplemented with 0.02% glycodiosgenin (GDN). The purity of the final protein was confirmed by SDS-PAGE. The protein concentration was determined by BCA Assay (Pierce) and used for biochemical assays or cryo-EM grid freezing.

### CryoEM sample prep and data acquisition

4 µl of concentrated NgFks1 (3 mg/ml) or in complex with caspofungin was applied to a QF-1.2/1.3-4Au 400 mesh grid and flash-frozen using a Leica EM GP2 equilibrated to 10 °C and 80% humidity. After a 10-second incubation, the sample was blotted for 4 seconds and then flash-frozen in liquid ethane.

For NgFks1 apo, a total of 5294 movies were captured on a Titan Krios at 300 kV equipped with a K3 Summit detector (Gatan) at the Pacific Northwest Center for Cryo-EM. The magnification was set to 130,000× in the automated super-resolution mode, and the defocus range was -0.8 to - 2.2 μm. The movies contained 62 frames with a total exposure/dose of 50 e⁻/Å².

For caspofungin-bound NgFks1, the purified NgFks1(3 mg/ml) was incubated with 200 μM caspofungin for 1 hr at 20 °C. The data was collected in EER mode on a Titan Krios at 300 kV equipped with a Falcon4i detector (Thermo). A total of 10,514 movies were collected. The magnification was 165,000x in non-super-resolution mode and the defocus range was -0.6 to 2.4 μm. The dose rate was exposure/dose of 32 e⁻/Å².

### Cryo-EM data processing

Both datasets were processed using CryoSPARC (version 4.7 and 5.0) ^47,48^. For the NgFks1 apo dataset, the imported movies were motion-corrected using Patch Motion Correction and the Contrast Transfer Function (CTF) was estimated using patch CTF. After denoising the micrographs, particle picking was performed using the blob picker. After multiple rounds of 2D classification, the best classes were selected to generate an ab initio reconstruction using 2 rounds of High-resolution ab initio reconstruction (HR-HAIR) parameters reported in a recent preprint ^49^. After rounds of local refinement, nonuniform refinement, global and local CTF refinement, and reference-based motion correction and ab initio refinement, the final reconstruction generated a structure of a 3.47 Å resolution cryo-EM map of NgFks1, which consists of 33,592 particles (Fig. S9).

For caspofungin-bound NgFks1, the movies were motion-corrected, and CTF parameters were estimated as mentioned before. Micrographs were filtered and processed using curate exposure followed by denoising. Initially, we took a small subset of the micrographs on which we picked particles using Blob Picker, generated 2D classes, and then, through an ab initio construction job, created a 3D volume. We used the generated volume as a template to pick particles from all the denoised micrographs. The selected particles were then 2D-classified, and we ended up with good classes composed of 82,712 particles. These 2D-classified particles were then processed using HR-HAIR parameters into three classes. The best class was then refined by rounds of local refinement, Global CTF refinement, Local CTF refinement, and non-uniform refinement, yielding a 3.24 Å map with 75,499 particles. In parallel, excluded particles from an earlier 2D classification were subjected to 5 ab initio reconstruction classes. The selected and excluded particles from the earlier 2D classification were combined, and a heterogeneous classification was performed using 5 ab initio maps, two excluded ab initio maps from the HR-HAIR job, and 3.24 Å maps as references. The best class consisting of 255,005 particles was combined with the 75,499 particles, followed by the removal of the duplicate particles. This particle set was further processed by non-uniform refinement, re-extraction/upsampling to the original pixel size, reference-based motion correction, and local refinement, producing a 2.98 Å reconstruction from 244,703 particles with an intact lipid tail but broken headgroup density. Following a round of 3D classification, the new ab initio refinement job was used in CryoSPARC5 to maintain gold standard FSC criteria with an entirely new map, with a re-splitting of particle half sets. Another round of 3D classification was performed to improve the drug density, followed by a final non-uniform refinement that generated a reconstruction at 3.32 Å estimated using the gold standard FSC with 112725 particles (Fig. S10).

### Model building and refinement

An initial Fks1 model was built using Boltz 2 ^50^. *Coot* version 0.9.8.93 ^51^ and Isolde 1.10.1 and 1.8 ^52^ were used in ChimeraX 1.8 and 1.10.1 ^53^ to fit the model into the cryo-EM density. Multiple cycles of real-space refinement analysis were performed in Phenix version 1.21.2 ^54^. The final figures were generated using ChimeraX 1.10.1. The restraints for caspofungin and glucan were generated using eLBOW in Phenix version 1.21.2 ^55^. During the Isolde simulation, residue parameterization for CPG was performed as implemented in Isolde. The final model refinement statistics for both models are shown in Table S1-2.

### Microscale thermophoresis assay

The 0.02% GDN purified protein was fixed at 200 nM using microBCA. Then the protein was incubated with a cysteine labeling dye at a protein: dye ratio of 1:10 in the dark for 45 min at 4 °C. The free dye was removed from the protein by using a 100-kDa molecular-weight-cutoff centrifugal concentrator. The labeled protein was diluted with buffer and then concentrated, twice to ensure efficient removal of the free dye. 50 µL of the protein-ligand mixtures were prepared using a constant concentration of protein and varying concentrations of caspofungin. Three independent measurements were performed with caspofungin concentrations ranging from 10 µM to 1600 µM, with a no-ligand control. Finally, the MST experiments were performed in a Monolith NT.115 pico instrument. Standard MST capillaries were used in the MST measurement.

### Cell survivability assay

The knockout fks1 strain (ΔFks1) was created using the homologous recombination method in the BJ2168 strain. The Fks1 mutants were generated using PCR-based mutagenesis. The constructs were transformed into the ΔFks1 strain and grown for 7 days on YNB-Ura plates. A single colony was picked and cultured at 30°C in the YNB-Ura medium and fixed to an OD600 of 1.0. 200 µL of this culture was then inoculated into 10 ml YRG-Ura (yeast nitrogen base with ammonium sulfate 0.67% wt/vol, raffinose 1% wt/vol, galactose 2% wt/vol, CSM-Ura 0.077% wt/vol) media containing multiple dilutions of CFN. After 24 hours of incubation, OD600 was measured. OD values were then normalized to the no-drug concentration. Three biological replicates were performed to generate the mean for the reported values.

## Data availability

The coordinates are deposited in the PDB with the accession code 11kd for NgFks1 Apo and 35ut NgFks1 CFN. The EM data are available in the EMDB with accession code EMD-75764 for NgFks1 Apo and EMD-77198 for NgFks1 CFN. (Both PDB and EMDB is now held for publication)

## Acknowledgements

This research was funded through a grant from the National Institute of Allergy and Infectious Diseases (NIAID) under grant number NIH R01 AI156270 (awarded to TM Tomasiak). The National NIH-Funded Pacific Northwest Center for Cryo-EM (PNCC) at Oregon Health and Science University (OHSU) funded a portion of the study under the proposal ID#160652. We acknowledge Nancy Meyer for her invaluable assistance in collecting the NgFks1 apo dataset and Dewight Williams at Arizona State University for collecting the NgFks1 CFN dataset.

## Author Contribution

Md Shamiul Islam Rasel: Overall project administration, Fks1 expression and purification, Cryo-EM sample prep, Cryo-EM data curation, data analysis, model building, validation, conducting yeast survival assay, microscale thermophoresis assay, and writing the manuscript.

Darpan Raghav: Construct design, Fks1 knockout strain generation, survival assay replicate run and reviewing and editing the manuscript.

Fahrin Jahan Suchana: Protein expression and purification.

R Souza Amado de Carvalho: Cryo-EM sample preparation and model building.

TM Tomasiak: Conceptualization, supervision, funding acquisition, validation, model building, writing, reviewing and editing the manuscript.

