## Supporting Information for "Structural basis of caspofungin inhibition of *Nakaseomyces glabratus* Fks1"

### Structural basis of caspofungin inhibition in the post-catalytic state of *Nakaseomyces glabratus* Fks1

**This PDF file includes:**

Figures S1 to S10
Tables S1 to S2
SI References

```
N.glabrata_Fks1     MSYNNNGQQ...............MSDQGYYDNNQGY.YQPED..QQNGQAM........
N.glabrata_Fks2     MSYDQGGNGNWQNTDPNGNYYYNGAENNEFYDQDYAS.QQPEQ..QQGGEGYYDEYGQPN
S.cerevisiae_Fks1   MNTDQQP.......YQGQTDYTQGPGN......GQ............SQEQDYDQYGQPL
S.cerevisiae_Fks2   MSYN.DP.......NLNGQYYSNGDGT......GDGN.YPTYQ..VTQDQSAYDEYGQPI
C.albicans_Fks1     MSYNDN....................NNHYYDPNQQGGMPPHQGGEGYYQQQYDDMGQQP
C.auris_Fks1        MSY.DN....................NHNYYDPNQQQGGQP..GGEYYQQGAYDEMGQPV

                                             40        50        60
N.glabrata_Fks1     ..........................YGDEGYYDPNISGGDYYNQ..PPPPPNMMGQDME
N.glabrata_Fks2     YNYMNDP.......QQGQMPQQQPGGYENDGYYDS......YYNNQMNAGVGNGLGPDQT
S.cerevisiae_Fks1   YPSQ..ADGYYDPNVAAGTEADMYGQQPPNESYDQDYTNGEYYGQ..PPNMA...AQDGE
S.cerevisiae_Fks2   YTQNQLDDGYYDPNEQYVDGTQFPQGQDPSQDQGPYNNDASYYNQ..PPNMMNPSSQDGE
C.albicans_Fks1     .....HQQDYYDPNAQYQQQPYDMDGYQDQANYGG............QPMNAQGYNADPE
C.auris_Fks1        NY..DQAGDYYDPNQQYQQQPYDMDGYQNYGQQ.................GAAGYSADPE

                      70        80        90        100       110
N.glabrata_Fks1     NFSDFSSYG..PPGVQGYDYQNMNQSGQYTPSQMSYNGDP.NSGSSTPIYGNGMS.....
N.glabrata_Fks2     NFSDFSSYG..PPPFQNNQANYTPSQLSYSNNGMGSNGMN.MSGSSTPVYGN........
S.cerevisiae_Fks1   NFSDFSSYG..PPGTPGYDS....YGGQYTASQMSYGEPN.SSGTSTPIYG.........
S.cerevisiae_Fks2   NFSDFSSYG..PPSGTY.......PNDQYTPSQMSYPDQD.GSSGASTPYGNGVVNGNGQ
C.albicans_Fks1     AFSDFSYGG.QTPGTPGYD....QYGTQYTPSQMSYGGDPRSSGASTPIYGG.....QGQ
C.auris_Fks1        AFSDFSYGGGHAPGTPGYD....QYGAQYTPSQMSYAGA.RSSGASTPIYGA......NP

                    120       130       140       150       160       170
N.glabrata_Fks1     AYDPNAIAMALPNDPYPAWTADSQSPVPIEQIEDVFIDLTNKFGFQRDSMRNMFDHFMTL
N.glabrata_Fks2     .YDPNAIAMTLPNDPYPAWTADPQSPVSIEQIEDVFIDLTNKFGFQRDSMRNIFDLFMTL
S.cerevisiae_Fks1   NYDPNAIAMALPNEPYPAWTADSQSPVSIEQIEDIFIDLTNRLGFQRDSMRNMFDHFMVL
S.cerevisiae_Fks2   YYDPNAIEMALPNDPYPAWTADPQSPLPIEQIEDIFIDLTNKFGFQRDSMRNMFDHFMTL
C.albicans_Fks1     GYDPTQFNMS.SNLPYPAWSADPQAPIKIEHIEDIFIDLTNKFGFQRDSMRNMFDYFMTL
C.auris_Fks1        NYDPSQFQLS.SNLPYPAWSADPQAPIKIEHIEDIFIDLTNKFGFQRDSMRNMFDYFMTL

                    180       190       200       210       220       230
N.glabrata_Fks1     LDSRTSRMAPDQALLSLHADYIGGDTANYKKWYFAAQLDMDDEVGFRNMSLGKLSRKARK
N.glabrata_Fks2     LDSRTSRMSPDQALLSVHADYIGGDTANYKKWYFAAQLDMDDEVGFRNMNLGKLSRKARK
S.cerevisiae_Fks1   LDSRSSRMSPDQALLSLHADYIGGDTANYKKWYFAAQLDMDDEIGFRNMSLGKLSRKARK
S.cerevisiae_Fks2   LDSRSSRMSPEQALLSLHADYIGGDTANYKKWYFAAQLDMDDEIGFRNMKLGKLSRKARK
C.albicans_Fks1     LDSRSSRMSPAQALLSLHADYIGGDNANYRKWYFSSQQDLDDSLGFANMTLGKIGRKARK
C.auris_Fks1        LDSRSSRMSPAQALLSLHADYIGGDNANYKKWFFASQQDLDESIGFANMNLGKIGRKARK

                    240       250       260       270       280       290
N.glabrata_Fks1     AKKKNKKAMEEA..NPEDAEDILNKLEGDNSLEAADFRWKTKMNALTPIERVRQIALYLL
N.glabrata_Fks2     AKKKNKKAMEEA..NPEDAAEVLNKIEGDNSLEASDFRWKTKMNMLTPIERVRQVALYML
S.cerevisiae_Fks1   AKKKNKKAMEEA..NPEDTEETLNKIEGDNSLEAADFRWKAKMNQLSPLERVRHIALYLL
S.cerevisiae_Fks2   AKKKNKKAMQEA..SPEDTEETLNQIEGDNSLEAADFRWKSKMNQLSPFEMVRQIALFLL
C.albicans_Fks1     ASKKSKKARKAAEEHGQDVDALANELEGDYSLEAAEIRWKAKMNSLTPEERVRDLALYLL
C.auris_Fks1        ASKKSKKARKAAEEHGQDIDALNNELEGDYSMEAANIRWKAKMNVLTPEERVRDIALYLL

                      300       310       320       330       340       350
N.glabrata_Fks1     IWGEANQVRFTSECLCFIYKCATDYLNSPLCQQRTEPMPEGDYLNRVITPLYRFIRNQVY
N.glabrata_Fks2     IWGEANQVRFTSECLCFIYKCASDYLESPLCQQRTEPIPEGDYLNRVITPIYQFIRNQVY
S.cerevisiae_Fks1   CWGEANQVRFTAECLCFIYKCALDYLDSPLCQQRQEPMPEGDFLNRVITPIYHFIRNQVY
S.cerevisiae_Fks2   CWGEANQVRFTPECLCFIYKCASDYLDSAQCQQRPDPLPEGDFLNRVITPLYRFIRSQVY
C.albicans_Fks1     IWGEANQVRFTPECLCYIYKSATDYLNSPLCQQRQEPVPEGDYLNRVITPLYRFIRSQVY
C.auris_Fks1        LWGEANQVRFTPELICFIFKTALDYLNSPQCQQRQEPVPEGDYLNRIITPIYRFIRSQVY
```

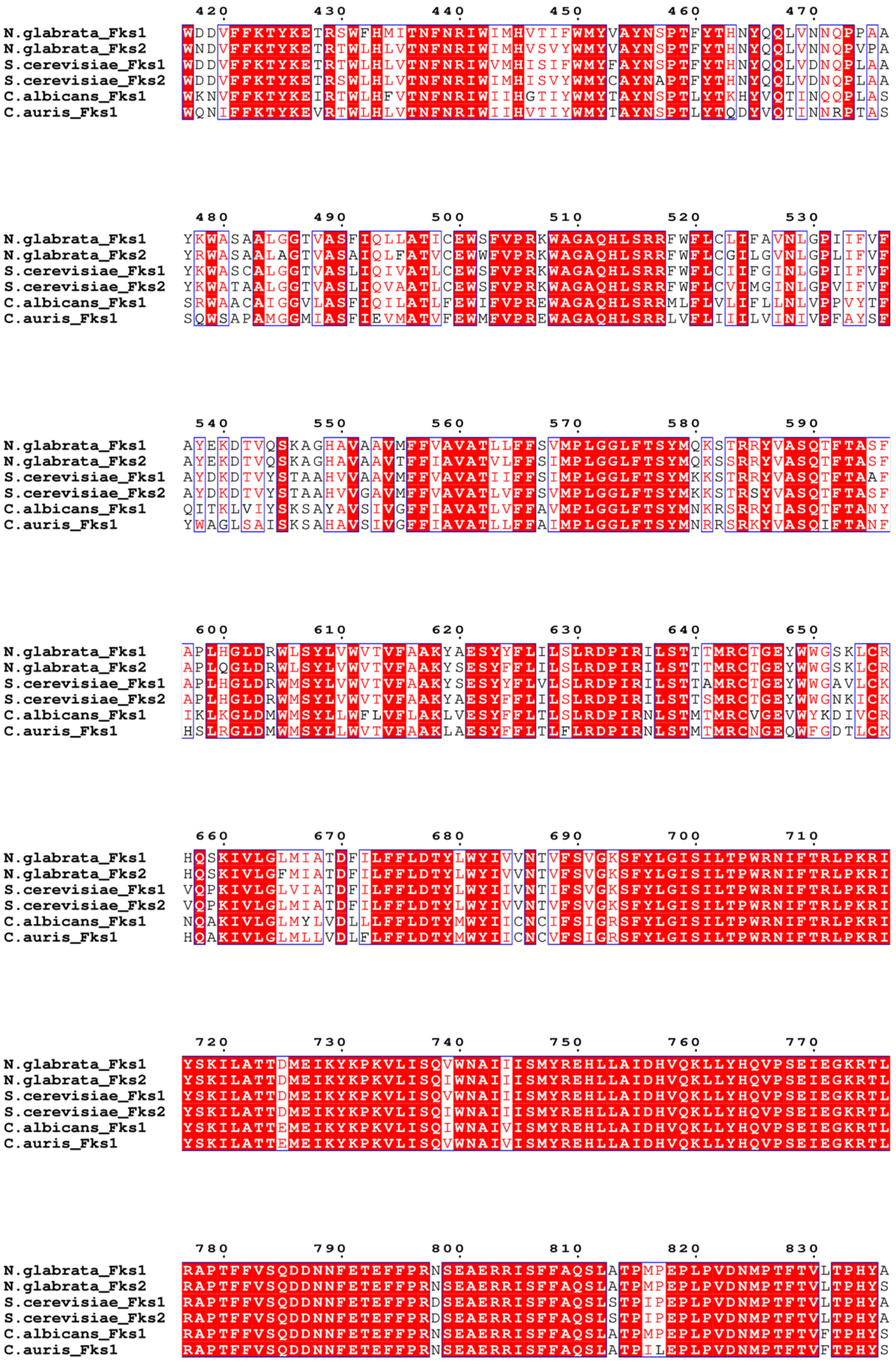
420 430 440 450 460 470
N.glabrata_Fks1 WDDVFFKTYKETRSWFHMITNFNRIWIMHVTIFWMYVAYNSPTFYTHNYQQLVNNQPPAA
N.glabrata_Fks2 WNDVFFKTYKETRTWLHLVTNFNRIWIMHVSVYWMYVAYNSPTFYTHNYQQLVNNQPVPA
S.cerevisiae_Fks1 WDDVFFKTYKETRTWLHLVTNFNRIWVMHISIFWMYFAYNSPTFYTHNYQQLVDNQPLAA
S.cerevisiae_Fks2 WDDVFFKTYKETRSWLHLVTNFNRIWIMHISVYWMYCAYNAPTFYTHNYQQLVDNQPLAA
C.albicans_Fks1 WKNVFFKTYKEIRTWLHFVTNFNRIWIIHGTIYWMYTAYNSPTLYTKHYVQTINQQPLAS
C.auris_Fks1 WQNIFFKTYKEVRTWLHLVTNFNRIWIIHVTIYWMYTAYNSPTLYTQDYVQTINNRPTAS
480 490 500 510 520 530
N.glabrata_Fks1 YKWASAALGGTVASFIQLLATICEWSFVPRKWAGAQHLSRRFWFLCLIFAVNLGPIIFVF
N.glabrata_Fks2 YRWASAALAGTVASAIQLFATVCEWWFVPRKWAGAQHLSRRFWFLCGILGVNLGPLIFVF
S.cerevisiae_Fks1 YKWASCALGGTVASLIQIVATLCEWSFVPRKWAGAQHLSRRFWFLCIIFGINLGPIIFVF
S.cerevisiae_Fks2 YKWATAALGGTVASLIQVAATLCEWSFVPRKWAGAQHLSRRFWFLCVIMGINLGPVIFVF
C.albicans_Fks1 SRWAACAIGGVLASFIQILATLFEWIFVPREWAGAQHLSRRMLFLVLIFLLNLVPPVYTF
C.auris_Fks1 SQWSAPAMGGMIASFIEVMATVFEWMFVPREWAGAQHLSRRLVFLIIILVINIVPFAYSF
540 550 560 570 580 590
N.glabrata_Fks1 AYEKDTVQSKAGHAVAAVMFFVAVATLLFFSVMPLGGLFTSYMQKSTRRYVASQTFTASF
N.glabrata_Fks2 AYEKDTVQSKAGHAVAAVTFFIAVATVLFFSIMPLGGLFTSYMQKSSRRYVASQTFTASF
S.cerevisiae_Fks1 AYDKDTVYSTAAHVVAAVMFFVAVATIIFFSIMPLGGLFTSYMKKSTRRYVASQTFTAAF
S.cerevisiae_Fks2 AYDKDTVYSTAAHVVGAVMFFVAVATLVFFSVMPLGGLFTSYMKKSTRSYVASQTFTASF
C.albicans_Fks1 QITKLVIYSKSAYAVSIVGFFIAVATLVFFAVMPLGGLFTSYMNKRSRRYIASQTFTANY
C.auris_Fks1 YWAGLSAISKSAHAVSIVGFFIAVATLLFFAIMPLGGLFTSYMNRRSRKYVASQIFTANF
600 610 620 630 640 650
N.glabrata_Fks1 APLHGLDRWLSYLVWVTVFAAKYAESYYFLILSLRDPIRILSTTTMRCTGEYWWGSKLCR
N.glabrata_Fks2 APLQGLDRWLSYLVWVTVFAAKYSESYYFLILSLRDPIRILSTTTMRCTGEYWWGSKLCR
S.cerevisiae_Fks1 APLHGLDRWMSYLVWVTVFAAKYSESYYFLVLSLRDPIRILSTTAMRCTGEYWWGAVLCK
S.cerevisiae_Fks2 APLHGLDRWMSYLVWVTVFAAKYAESYFFLILSLRDPIRILSTTSMRCTGEYWWGNKICK
C.albicans_Fks1 IKLKGLDMWMSYLLWFLVFLAKLVESYFFLTLSLRDPIRNLSTMTMRCVGEVWYKDIVCR
C.auris_Fks1 HSLRGLDMWMSYLLWVTVFAAKLAESYFFLTLFLRDPIRNLSTMTMRCNGEQWFGDTLCK
660 670 680 690 700 710
N.glabrata_Fks1 HQSKIVLGLMIATDFILFFLDTYLWYIVVNTVFSVGKSFYLGISILTPWRNIFTRLPKRI
N.glabrata_Fks2 HQSKIVLGFMIATDFILFFLDTYLWYIVVNTVFSVGKSFYLGISILTPWRNIFTRLPKRI
S.cerevisiae_Fks1 VQPKIVLGLVIATDFILFFLDTYLWYIIVNTIFSVGKSFYLGISILTPWRNIFTRLPKRI
S.cerevisiae_Fks2 VQPKIVLGLMIATDFILFFLDTYLWYIVVNTVFSVGKSFYLGISILTPWRNIFTRLPKRI
C.albicans_Fks1 NQAKIVLGLMYLVDLLLFFLDTYMWYIICNCIFSIGRSFYLGISILTPWRNIFTRLPKRI
C.auris_Fks1 HQAKIVLGLMLLVDLFLFFLDTYMWYIICNCVFSIGRSFYLGISILTPWRNIFTRLPKRI
720 730 740 750 760 770
N.glabrata_Fks1 YSKILATTDMEIKYKPKVLISQVWNAIIISMYREHLLAIDHVQKLLYHQVPSEIEGKRTL
N.glabrata_Fks2 YSKILATTDMEIKYKPKVLISQIWNAIIISMYREHLLAIDHVQKLLYHQVPSEIEGKRTL
S.cerevisiae_Fks1 YSKILATTDMEIKYKPKVLISQVWNAIIISMYREHLLAIDHVQKLLYHQVPSEIEGKRTL
S.cerevisiae_Fks2 YSKILATTDMEIKYKPKVLISQIWNAIIISMYREHLLAIDHVQKLLYHQVPSEIEGKRTL
C.albicans_Fks1 YSKILATTEMEIKYKPKVLISQIWNAIVISMYREHLLAIDHVQKLLYHQVPSEIEGKRTL
C.auris_Fks1 YSKILATTEMEIKYKPKVLISQVWNAIVISMYREHLLAIDHVQKLLYHQVPSEIEGKRTL
780 790 800 810 820 830
N.glabrata_Fks1 RAPTFFVSQDDNNFETEFFPRNSEAERRISFFAQSLATPMPEPLPVDNMPTFTVLTPHYA
N.glabrata_Fks2 RAPTFFVSQDDNNFETEFFPRNSEAERRISFFAQSLATPMPEPLPVDNMPTFTVLTPHYS
S.cerevisiae_Fks1 RAPTFFVSQDDNNFETEFFPRDSEAERRISFFAQSLSTPIPEPLPVDNMPTFTVLTPHYA
S.cerevisiae_Fks2 RAPTFFVSQDDNNFETEFFPRDSEAERRISFFAQSLSTPIPEPLPVDNMPTFTVLTPHYA
C.albicans_Fks1 RAPTFFVSQDDNNFETEFFPRNSEAERRISFFAQSLATPMPEPLPVDNMPTFTVFTPHYS
C.auris_Fks1 RAPTFFVSQDDNNFETEFFPRNSEAERRISFFAQSLATPILEPLPVDNMPTFTVFTPHYS

```
                   900       910       920       930       940       950
N.glabrata_Fks1    KEDALKNQIDDLPFYCIGFKSAAPEYTLRTRIWASLRSQTLYRTVSGFMNYARAIKLLYR
N.glabrata_Fks2    KSDALKTQIDDLPFYCIGFKSAAPEYTLRTRIWASLRSQTLYRTVSGFMNYARAIKLLYR
S.cerevisiae_Fks1  KEDALKSQIDDLPFYCIGFKSAAPEYTLRTRIWASLRSQTLYRTISGFMNYSRAIKLLYR
S.cerevisiae_Fks2  KEDALKSQIDDLPFYCIGFKSAAPEYTLRTRIWASLRSQTLYRTISGFMNYSRAIKLLYR
C.albicans_Fks1    SEDGLKSKIDDLPFYCIGFKSAAPEYTLRTRIWASLRSQTLYRTVSGFMNYARAIKLLYR
C.auris_Fks1       NEDGLKAKIDDLPFYCIGFKSAAPEYTLRTRIWASLRSQTLYRTVSGFMNYARAIKLLYR

                   960       970       980       990       1000      1010
N.glabrata_Fks1    VENPEIVQMFGGNAEGLERELEKMARRKFKFLVSMQRLAKFKPHELENAEFLLRAYPDLQ
N.glabrata_Fks2    VENPEIVQMFGGNAEGLERELEKMARRKFKFLVSMQRLAKFKPHELENTEFLLRAYPDLQ
S.cerevisiae_Fks1  VENPEIVQMFGGNAEGLERELEKMARRKFKFLVSMQRLAKFKPHELENAEFLLRAYPDLQ
S.cerevisiae_Fks2  VENPEIVQMFGGNADGLERELEKMARRKFKFLVSMQRLAKFKPHELENAEFLLRAYPDLQ
C.albicans_Fks1    VENPELVQYFGGDPEGLELALERMARRKFRFLVSMQRLSKFKDDEMENAEFLLRAYPDLQ
C.auris_Fks1       VENPELVQYFGGDPEGLELALEKMARRKFKFVVSMQRLAKFKEDEMENAEFLLRAYPDLQ

                   1020      1030      1040      1050      1060      1070
N.glabrata_Fks1    IAYLDEEPPLNEGEEPRIYSALIDGHCEILENGRRRPKFRVQLSGNPILGDGKSDNQNHA
N.glabrata_Fks2    IAYLDEEPPLNEGEEPRIYSALIDGHCEMLENGRRRPKFRVQLSGNPILGDGKSDNQNHA
S.cerevisiae_Fks1  IAYLDEEPPLTEGEEPRIYSALIDGHCEILDNGRRRPKFRVQLSGNPILGDGKSDNQNHA
S.cerevisiae_Fks2  IAYLDEEPPLNEGEEPRIYSALIDGHCEILENGRRRPKFRVQLSGNPILGDGKSDNQNHA
C.albicans_Fks1    IAYLDEEPALNEDEEPRVYSALIDGHCEMLENGRRRPKFRVQLSGNPILGDGKSDNQNHA
C.auris_Fks1       IAYLDEEPPLNEDEEPRVYSALIDGHCEVLDNGRRRPKFRVQLSGNPILGDGKSDNQNHA

                   1080      1090      1100      1110      1120      1130
N.glabrata_Fks1    LIFYRGEYIQLIDANQDNYLEECLKIRSVLAEFEELNAEQVYPYSPGVKYEDQN.TNHPV
N.glabrata_Fks2    LIFYRGEYIQLIDANQDNYLEECLKIRSVLAEFEELNAEPVYPYTPGVKYEDQK.TNHPV
S.cerevisiae_Fks1  LIFYRGEYIQLIDANQDNYLEECLKIRSVLAEFEELNVEQVNPYAPGLRYEEQT.TNHPV
S.cerevisiae_Fks2  LIFYRGEYIQLIDANQDNYLEECLKIRSVLAEFEELGIEQIHPYTPGLKYEDQS.TNHPV
C.albicans_Fks1    VIFHRGEYIQLIDANQDNYLEECLKIRSVLAEFEEMNVEHVNPYAPNLKSEDNNTKKDPV
C.auris_Fks1       IIFHRGEYIQLIDANQDNYLEECLKIRSVLAEFEELNVEHVNPYAPGLKNN.NDEKPAPV

                   1140      1150      1160      1170      1180      1190
N.glabrata_Fks1    AIVGAREYIFSENSGVLGDVAAGKEQTFGTLFARTLAQIGGKLHYGHPDFINATFMTTRG
N.glabrata_Fks2    AIVGAREYIFSENSGVLGDVAAGKEQTFGTLFARTLAQIGGKLHYGHPDFINATFMTTRS
S.cerevisiae_Fks1  AIVGAREYIFSENSGVLGDVAAGKEQTFGTLFARTLSQIGGKLHYGHPDFINATFMTTRG
S.cerevisiae_Fks2  AIVGAREYIFSENSGVLGDVAAGKEQTFGTLFARTLAQIGGKLHYGHPDFINATFMTTRG
C.albicans_Fks1    AFLGAREYIFSENSGVLGDVAAGKEQTFGTLFARTLAQIGGKLHYGHPDFLNATFMLTRG
C.auris_Fks1       AILGAREYIFSENSGVLGDVAAGKEQTFGTLFARTLAQIGGKLHYGHPDFLNATFMLTRG

                   1200      1210      1220      1230      1240      1250
N.glabrata_Fks1    GISKAQKGLHLNEDIYAGMNALLRGGRIKHCEYYQCGKGRDLGFGTILNFTTKIGAGMGE
N.glabrata_Fks2    GLSKAQKGLHLNEDIYAGMNALLRGGRIKHCEYYQCGKGRDLGFGTILNFTTKIGAGMGE
S.cerevisiae_Fks1  GVSKAQKGLHLNEDIYAGMNAMLRGGRIKHCEYYQCGKGRDLGFGTILNFTTKIGAGMGE
S.cerevisiae_Fks2  GVSKAQKGLHLNEDIYAGMNAVLRGGRIKHCEYYQCGKGRDLGFGTILNFTTKIGAGMGE
C.albicans_Fks1    GVSKAQKGLHLNEDIYAGMNAMMRGGKIKHCEYYQCGKGRDLGFGSILNFTTKIGAGMGE
C.auris_Fks1       GVSKAQKGLHLNEDIYAGMTAMLRGGRIKHCEYYQCGKGRDMGFGSICNFTTKIGAGMGE

                   1260      1270      1280      1290      1300      1310
N.glabrata_Fks1    QMLSREYYYLGTQLPIDRFLTFYYAHPGFHLNNLFIQLSLQMFMLTLVNLHALAHESIIC
N.glabrata_Fks2    QMLSREYYYLGTQLPVDRFLTFYYAHPGFHLNNLFIQLSLQMFMLTLVNLHALAHESILC
S.cerevisiae_Fks1  QMLSREYYYLGTQLPVDRFLTFYYAHPGFHLNNLFIQLSLQMFMLTLVNLSSLAHESIMC
S.cerevisiae_Fks2  QMLSREYYYLGTQLPIDRFLTFYYAHPGFHLNNLFIQLSLQMFMLTLVNLHALAHESILC
C.albicans_Fks1    QMLSREYFYLGTQLPLDRFLSFYYGHPGFHINNLFIQLSLQVFILVLANLNSLAHEAIMC
C.auris_Fks1       QMLSREYYYLSTQLPLDRFLSFYYGHPGFHINNLFIQLSLQTFMIVLANLNSLAHESILC
```

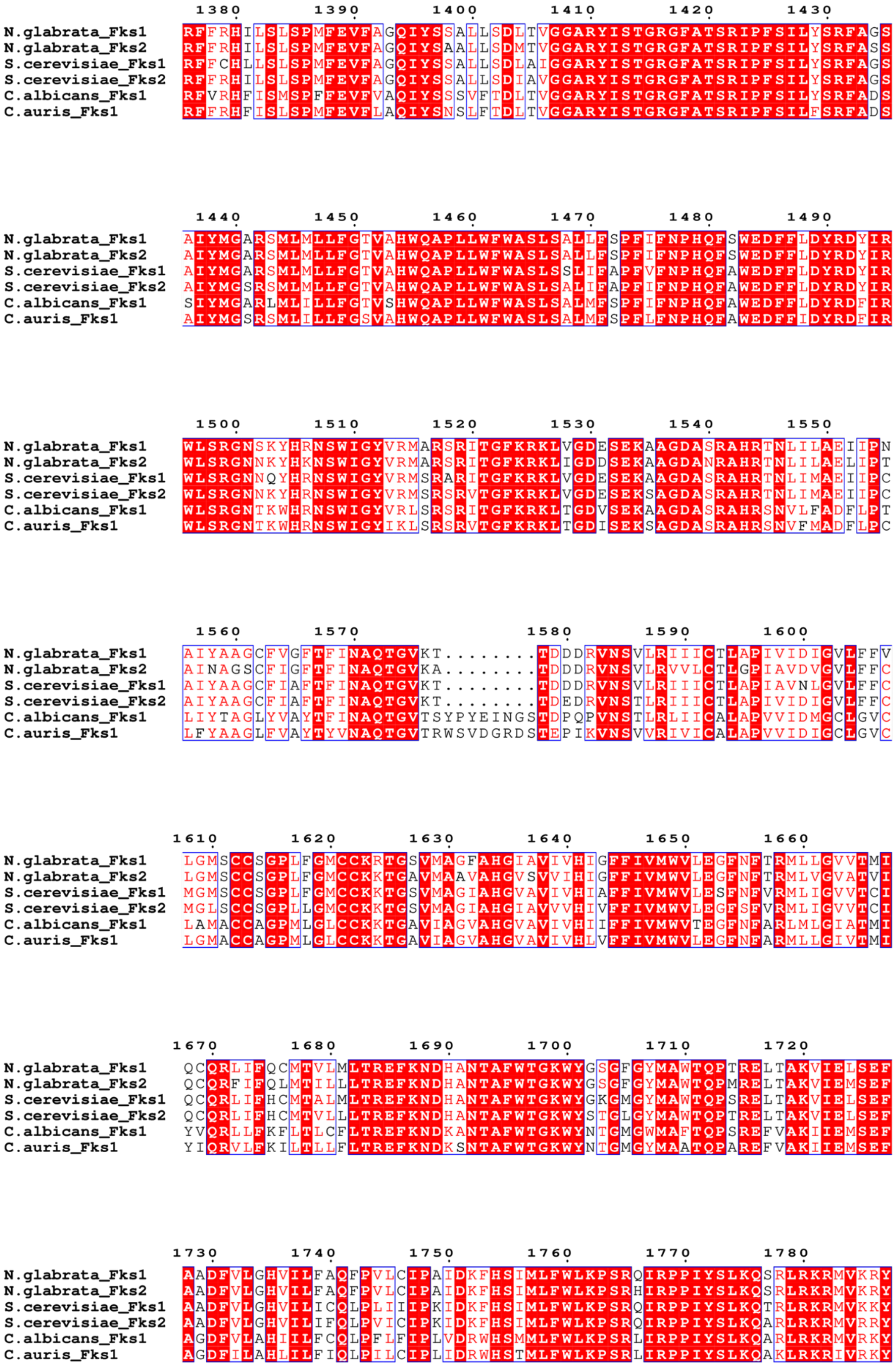
1380 1390 1400 1410 1420 1430
N.glabrata_Fks1 RFFRHILSLSPMFEVFAGQIYSSALLSDLTVGGARYISTGRGFATSRIPFSILYSRFAGS
N.glabrata_Fks2 RFFRHILSLSPMFEVFAGQIYSAALLSDMTVGGARYISTGRGFATSRIPFSILYSRFASS
S.cerevisiae_Fks1 RFFCHLLSLSPMFEVFAGQIYSSALLSDLAIGGARYISTGRGFATSRIPFSILYSRFAGS
S.cerevisiae_Fks2 RFFRHILSLSPMFEVFAGQIYSSALLSDIAVGGARYISTGRGFATSRIPFSILYSRFAGS
C.albicans_Fks1 RFVRHFISMSPFFEVFVAQIYSSSVFTDLTVGGARYISTGRGFATSRIPFSILYSRFADS
C.auris_Fks1 RFFRHFISLSPMFEVFLAQIYSNSLFTDLTVGGARYISTGRGFATSRIPFSILFSRFADS
1440 1450 1460 1470 1480 1490
N.glabrata_Fks1 AIYMGARSMLMLLFGTVAHWQAPLLWFWASLSALLFSPFIFNPHQFSWEDFFLDYRDYIR
N.glabrata_Fks2 AIYMGARSMLMLLFGTVAHWQAPLLWFWASLSALLFSPFIFNPHQFSWEDFFLDYRDYIR
S.cerevisiae_Fks1 AIYMGARSMLMLLFGTVAHWQAPLLWFWASLSSLIFAPFVFNPHQFAWEDFFLDYRDYIR
S.cerevisiae_Fks2 AIYMGSRSMLMLLFGTVAHWQAPLLWFWASLSALIFAPFIFNPHQFAWEDFFLDYRDYIR
C.albicans_Fks1 SIYMGARLMLILLFGTVSHWQAPLLWFWASLSALMFSPFIFNPHQFAWEDFFLDYRDFIR
C.auris_Fks1 AIYMGSRSMLILLFGSVAHWQAPLLWFWASLSALMFSPFLFNPHQFAWEDFFIDYRDFIR
1500 1510 1520 1530 1540 1550
N.glabrata_Fks1 WLSRGNSKYHRNSWIGYVRMARSRITGFKRKLVGDESEKAAGDASRAHRTNLILAEIIPN
N.glabrata_Fks2 WLSRGNNKYHKNSWIGYVRMARSRITGFKRKLIGDDSEKAAGDANRAHRTNLILAELIPT
S.cerevisiae_Fks1 WLSRGNNQYHRNSWIGYVRMSRARITGFKRKLVGDESEKAAGDASRAHRTNLIMAEIIPC
S.cerevisiae_Fks2 WLSRGNNKYHRNSWIGYVRMSRSRVTGFKRKLVGDESEKSAGDASRAHRTNLIMAEIIPC
C.albicans_Fks1 WLSRGNTKWHRNSWIGYVRLSRSRITGFKRKLTGDVSEKAAGDASRAHRSNVLFADFLPT
C.auris_Fks1 WLSRGNTKWHRNSWIGYIKLSRSRVTGFKRKLTGDISEKSAGDASRAHRSNVFMADFLPC
1560 1570 1580 1590 1600
N.glabrata_Fks1 AIYAAGCFVGFTFINAQTGVKT........TDDDRVNSVLRIIICTLAPIVIDIGVLFFV
N.glabrata_Fks2 AINAGSCFIGFTFINAQTGVKA........TDDDRVNSVLRVVLCTLGPIAVDVGVLFFC
S.cerevisiae_Fks1 AIYAAGCFIAFTFINAQTGVKT........TDDDRVNSVLRIIICTLAPIAVNLGVLFFC
S.cerevisiae_Fks2 AIYAAGCFIAFTFINAQTGVKT........TDEDRVNSTLRIIICTLAPIVIDIGVLFFC
C.albicans_Fks1 LIYTAGLYVAYTFINAQTGVTSYPYEINGSTDPQPVNSTLRLIICALAPVVIDMGCLGVC
C.auris_Fks1 LFYAAGLFVAYTYVNAQTGVTRWSVDGRDSTEPIKVNSVVRIVICALAPVVIDIGCLGVC
1610 1620 1630 1640 1650 1660
N.glabrata_Fks1 LGMSCCSGPLFGMCCKRTGSVMAGFAHGIAVIVHIGFFIVMWVLEGFNFTRMLLGVVTMI
N.glabrata_Fks2 LGMSCCSGPLFGMCCKKTGAVMAAVAHGVSVVIHIGFFIVMWVLEGFNFTRMLVGVATVI
S.cerevisiae_Fks1 MGMSCCSGPLFGMCCKKTGSVMAGIAHGVAVIVHIAFFIVMWVLESFNFVRMLIGVVTCI
S.cerevisiae_Fks2 MGLSCCSGPLLGMCCKKTGSVMAGIAHGIAVVVHIVFFIVMWVLEGFSFVRMLIGVVTCI
C.albicans_Fks1 LAMACCAGPMLGLCCKKTGAVIAGVAHGVAVIVHIIFFIVMWVTEGFNFARLMLGIATMI
C.auris_Fks1 LGMACCAGPMLGLCCKKTGAVIAGVAHGVAVIVHLVFFIVMWVLEGFNFARMLLGIVTMI
1670 1680 1690 1700 1710 1720
N.glabrata_Fks1 QCQRLIFQCMTVLMLTREFKNDHANTAFWTGKWYGSGFGYMAWTQPTRELTAKVIELSEF
N.glabrata_Fks2 QCQRFIFQLMTILLLTREFKNDHANTAFWTGKWYGSGFGYMAWTQPMRELTAKVIEMSEF
S.cerevisiae_Fks1 QCQRLIFHCMTALMLTREFKNDHANTAFWTGKWYGKGMGYMAWTQPSRELTAKVIELSEF
S.cerevisiae_Fks2 QCQRLIFHCMTVLLLTREFKNDHANTAFWTGKWYSTGLGYMAWTQPTRELTAKVIELSEF
C.albicans_Fks1 YVQRLLFKFLTLCFLTREFKNDKANTAFWTGKWYNTGMGWMAFTQPSREFVAKIIEMSEF
C.auris_Fks1 YIQRVLFKILTLLFLTREFKNDKSNTAFWTGKWYNTGMGYMAATQPAREFVAKIIEMSEF
1730 1740 1750 1760 1770 1780
N.glabrata_Fks1 AADFVLGHVILFAQFPVLCIPAIDKFHSIMLFWLKPSRQIRPPIYSLKQSRLRKRMVKRY
N.glabrata_Fks2 AADFVLGHVILFAQFPVLCIPAIDKFHSIMLFWLKPSRHIRPPIYSLKQSRLRKRMVKRY
S.cerevisiae_Fks1 AADFVLGHVILICQLPLIIIPKIDKFHSIMLFWLKPSRQIRPPIYSLKQTRLRKRMVKKY
S.cerevisiae_Fks2 AADFVLGHVILIFQLPVICIPKIDKFHSIMLFWLKPSRQIRPPIYSLKQARLRKRMVRRY
C.albicans_Fks1 AGDFVLAHIILFCQLPFLFIPLVDRWHSMMLFWLKPSRLIRPPIYSLKQARLRKRMVRKY
C.auris_Fks1 AGDFILAHLILFIQLPILCIPLIDRWHSTMLFWLKPSRLIRPPIYSLKQAKLRKRIVRKY

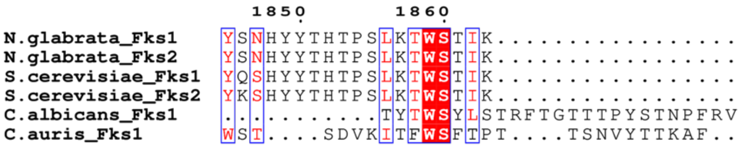

**Fig. S1. Multiple sequence alignment of Fks.** Fks1 and Fks2 sequences from N. glabrata (Uniprot: Q6FTN8, Q6FMZ6), S. cerevisiae (P38631, P40989 ), Fks1 C. auris (Uniprot: A0A3G2LW13), and the Fks1 sequence from C. Albicans (A0A1D8PCT0), are aligned using Clustal Omega (1)and then displayed using ESPript 3.2 (2).

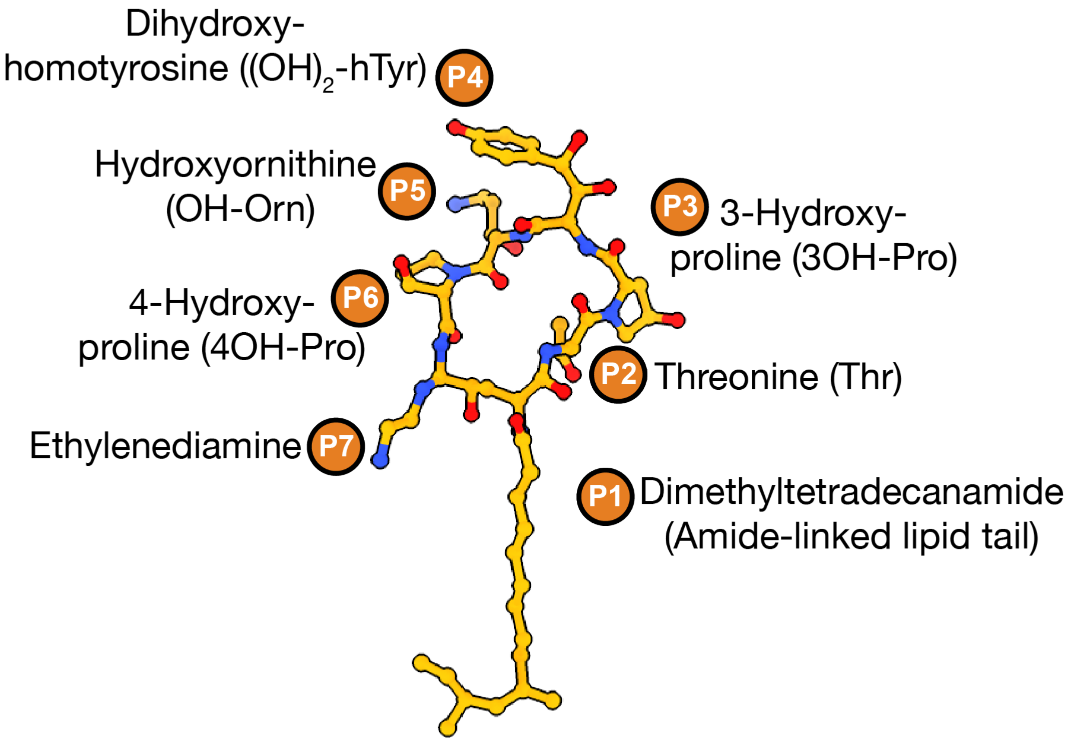

**Fig. S2. Structural moieties of caspofungin.** The lipidic tail is shown in a yellow shade containing the dimethyltetradecanamide (P1) group. The cyclic peptidic core is annotated in an orange shade, containing threonine (P2), 3-hydroxyproline (P3), dihydroxyhomotyrosine (P4), hydroxyornithine (P5), 4-hydroxyproline (P6), and the ethylenediamine (P7) group.

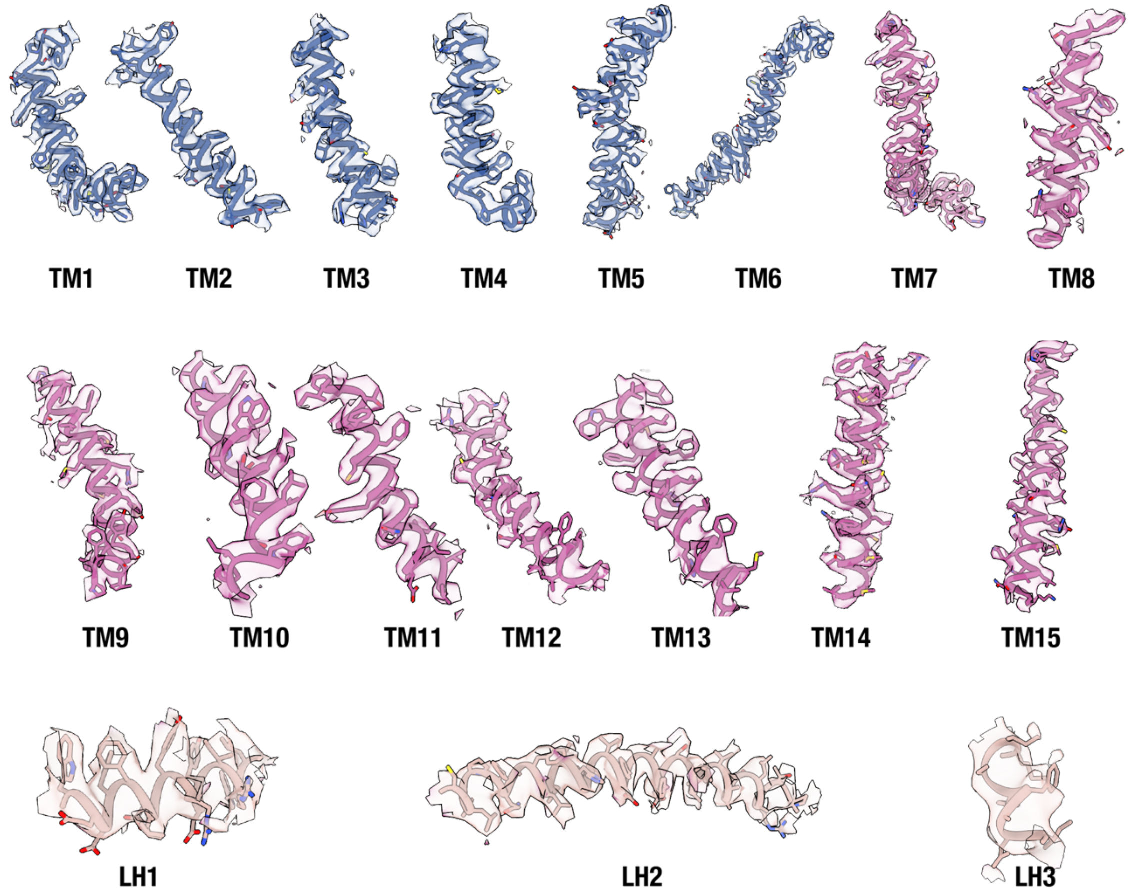

**Fig. S3A. TM helices with Cryo-EM density for NgFks1 apo.** Isolated TM helices and Lateral helices are shown along with their maps. TM1-6 is colored blue, TM7-15 is colored purple and Lateral helices are colored light tan. Maps shown are at a contour level of 0.075.

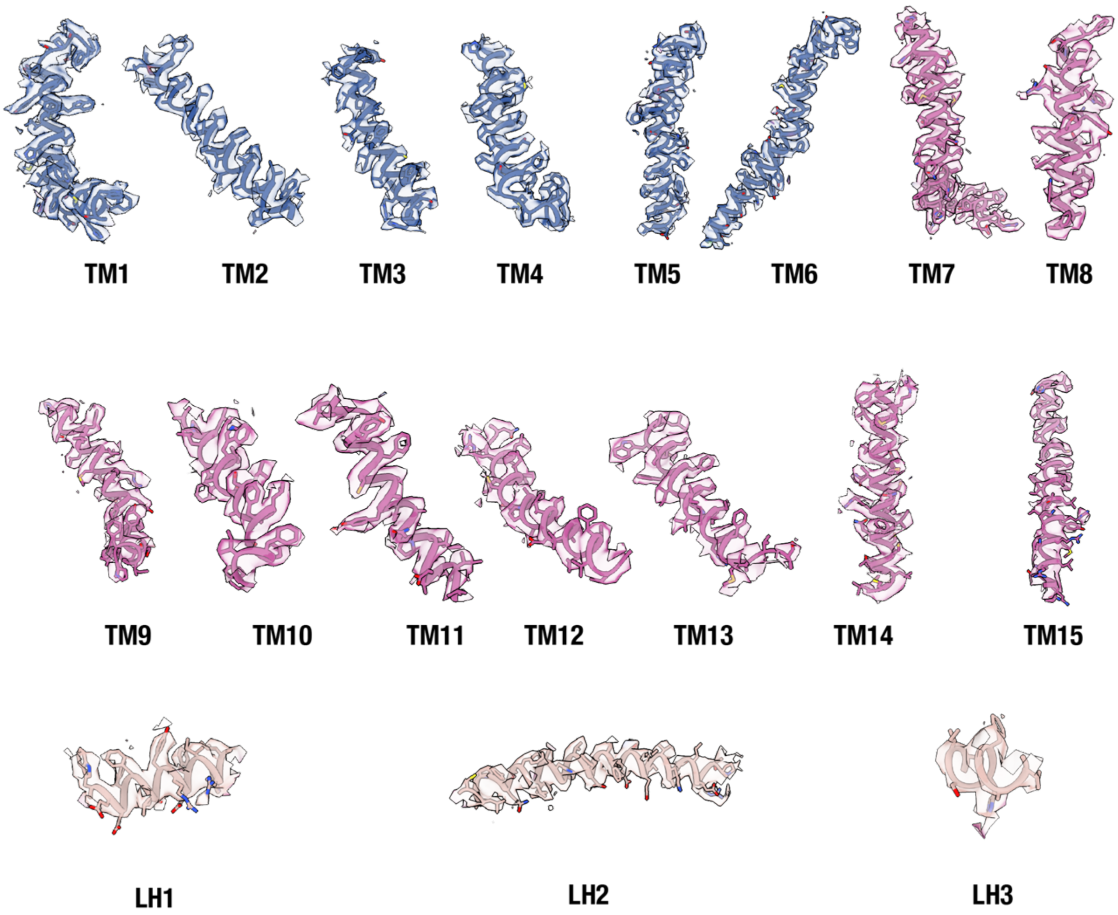

**Fig. S3B. TM helices with Cryo-EM density for NgFks1 CFN.** Isolated TM helices and Lateral helices are shown along with their maps. TM1-5 is colored blue, TM7-15 is colored purple and Lateral helices are colored light tan. Maps shown are at a contour level of 0.03.

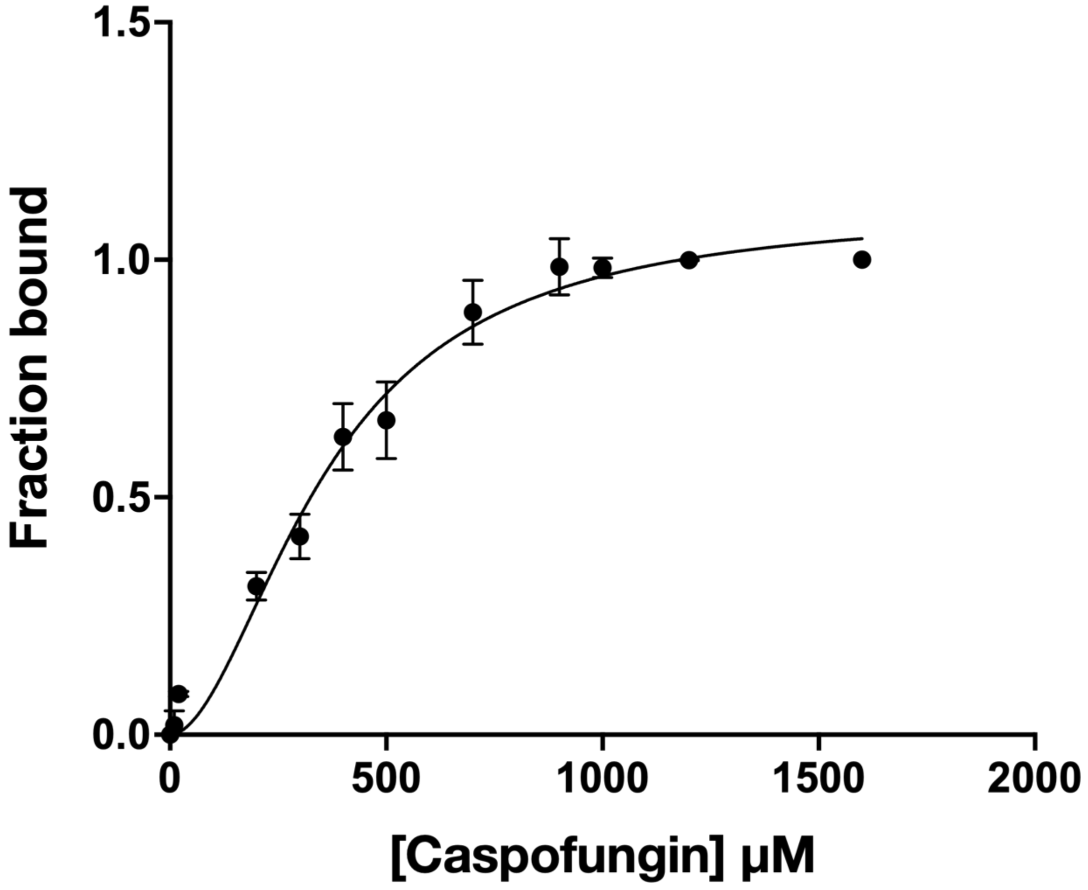

**Fig. S4. Microscale Thermophoresis assay for caspofungin to Fks1.** Fraction bound was plotted as a function of caspofungin concentration. Data were normalized from 0 (unbound) to 1 (fully bound). Error bars indicate variation among three replicate measurements where applicable.

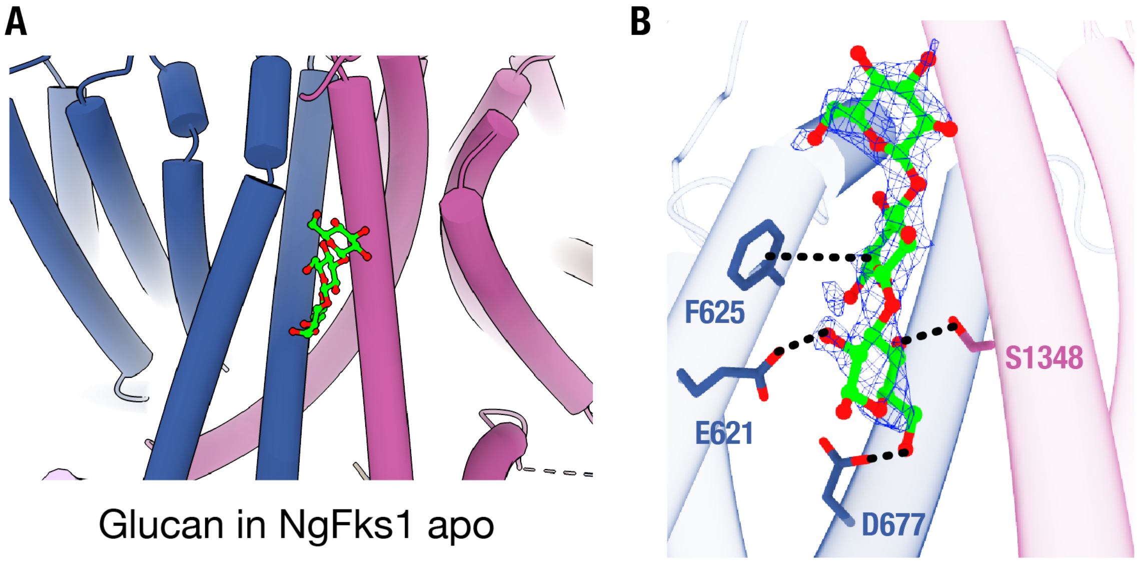

**Fig. S5. Glucan-bound NgFks1 apo structure. (A)** Overall view showing the position of the modeled glucan (lime) within the transmembrane helices of NgFks1. **(B)** Close-up view of the glucan model in the EM density shown in blue mesh with nearby stabilizing residues including E621, F625, D677 and S1348.

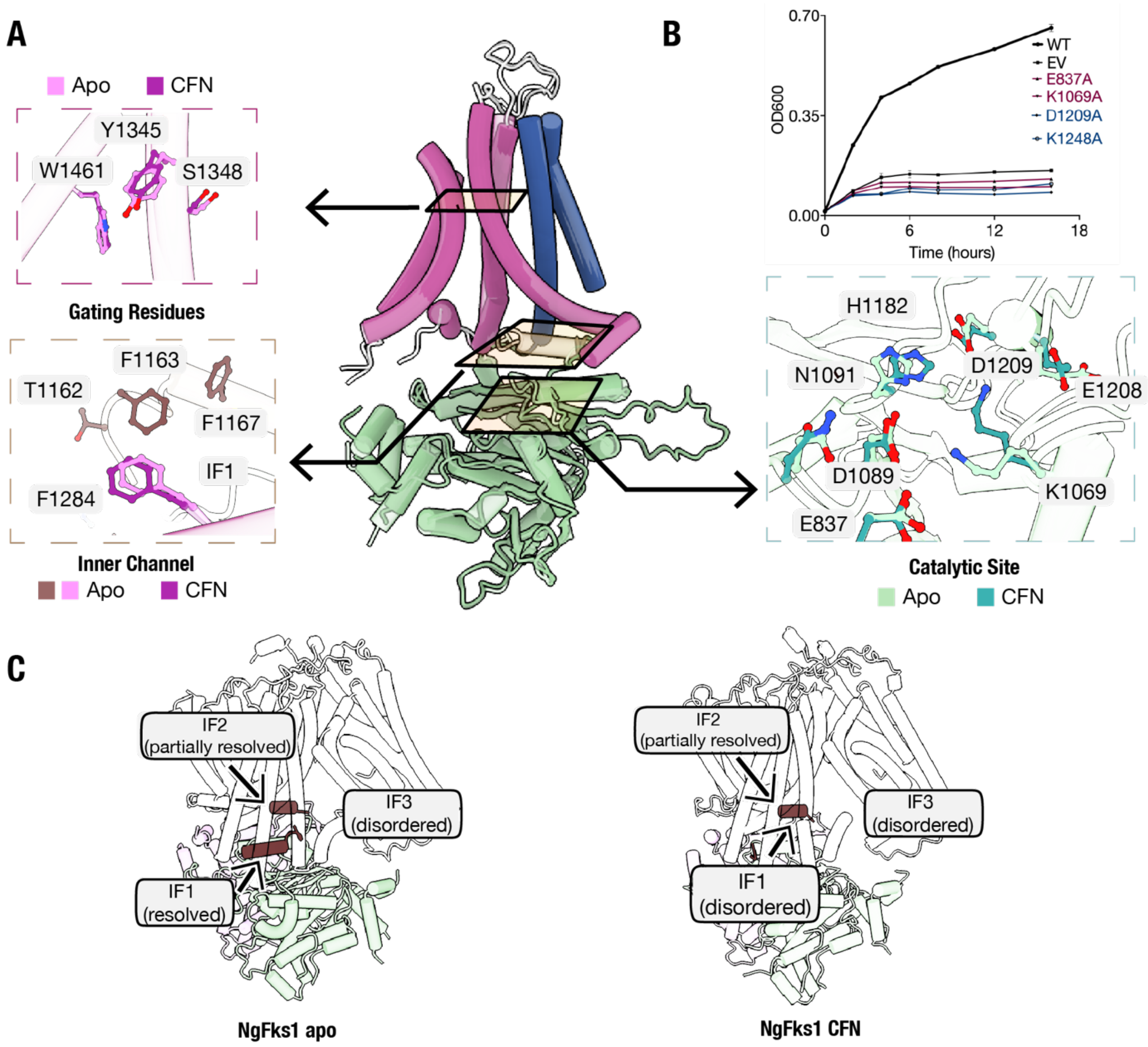

**Fig. S6. Conserved architecture of the glucan translocation path. (A)** The translocation path of NgFks1 based on the structure from pdb: 9pe1. The central channel is shown in the main view- the solid color represents the CFN bound state with the superimposition of the apo in transparent. The boxed inset shows a close-up comparison at key regions of the pathway: the upper exit gate (top left; CFN-bound in purple, apo in pink), the inner channel near IF1 (bottom left; CFN-bound in purple, apo in pink, with IF1 highlighted in dark brown), and the catalytic site (bottom right; CFN-bound in dark cyan, apo in light cyan). **(B)** Liquid growth assay of the catalytic site mutants. Each data point is a representation of 3 biological replicates. **(C)** Comparison of interfacial helices (IF1-IF3) organization in the apo and caspofungin-bound state. IF1 is resolved in apo but disordered in the CFN-bound state. IF2 is partially resolved and IF3 is disordered in both states.

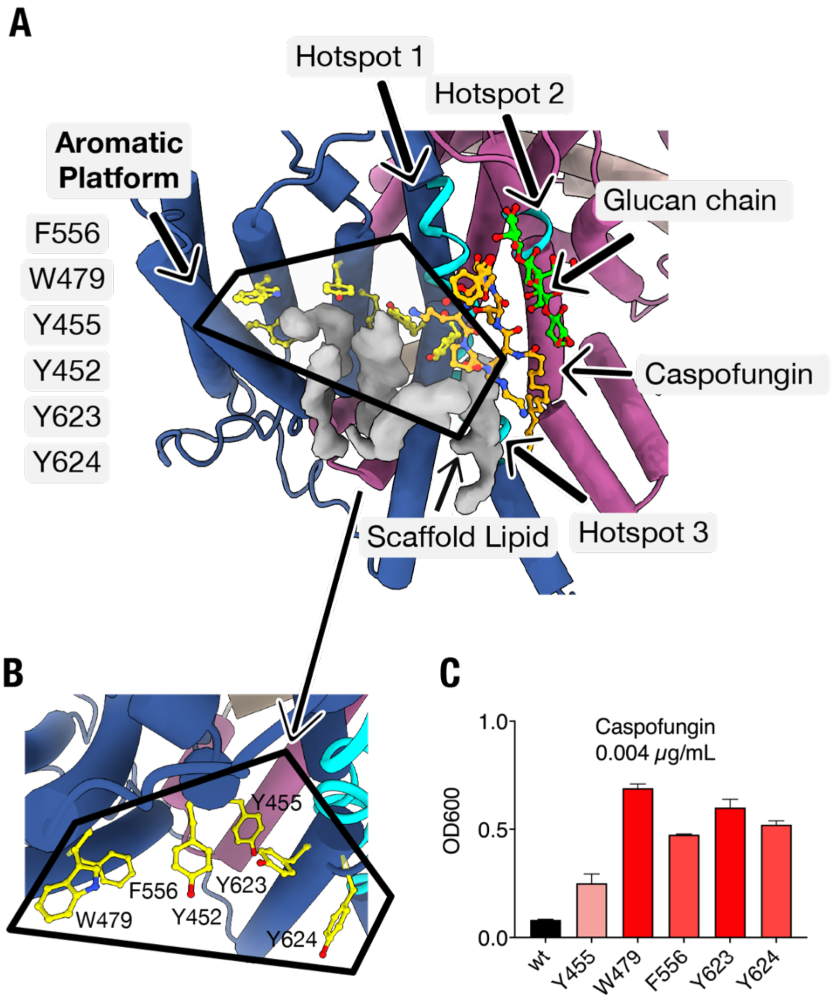

**Fig. S7. Hotspot 1 adjacent aromatic lipid pocket. (A)** Overview of the lipid densities (grey) relative to the glucan chain (lime), hotspot 1-3 (cyan), CFN (gold), and aromatic residues (yellow) (Y452, Y455, W479, F556, Y623, and Y624) surrounding the pocket like a platform. The lipid interacting with the caspofungin has been shown as a scaffold lipid **(B)** Close-up view of the aromatic platform residues. **(C)** Growth assay for aromatic platform alanine mutants in the presence of 0.004 µg/ml caspofungin showing a trend of reduction in caspofungin inhibition.

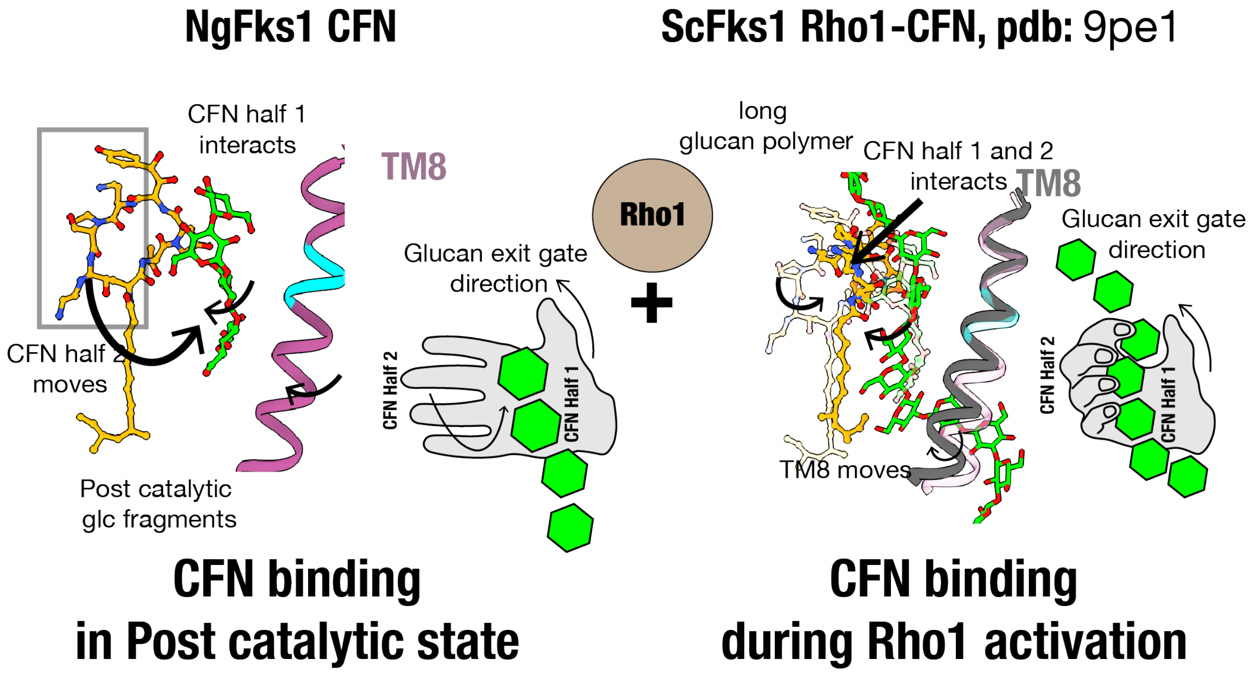

**Fig. S8. Proposed model for caspofungin engagement and glucan-dependent stabilization.** Comparison of our structure of the CFN-bound-post catalytic state of NgFks1 with the published structure of long glucan-CFN-bound ScFks1 in the rho1 activated state (pdb: 9pe1) (3). In our structure, CFN binds to Glc fragments on one side of the molecule (half1), while in the 9pe1 structure, both ends bind and capture the growing glucan chain. The cartoon representation by the side shows the mechanism, where the palm and figures represent half 1 and half 2 of the caspofungin molecule, respectively and the direction of the thumb represents the glucan exit gate. In this model, our structure is comparable to an open hand where the glucan fragment binds with the palm during the post-catalytic resting state. In the 9pe1 structure, the growing glucan chain is captured through a claw engaging both fingers and the palm.

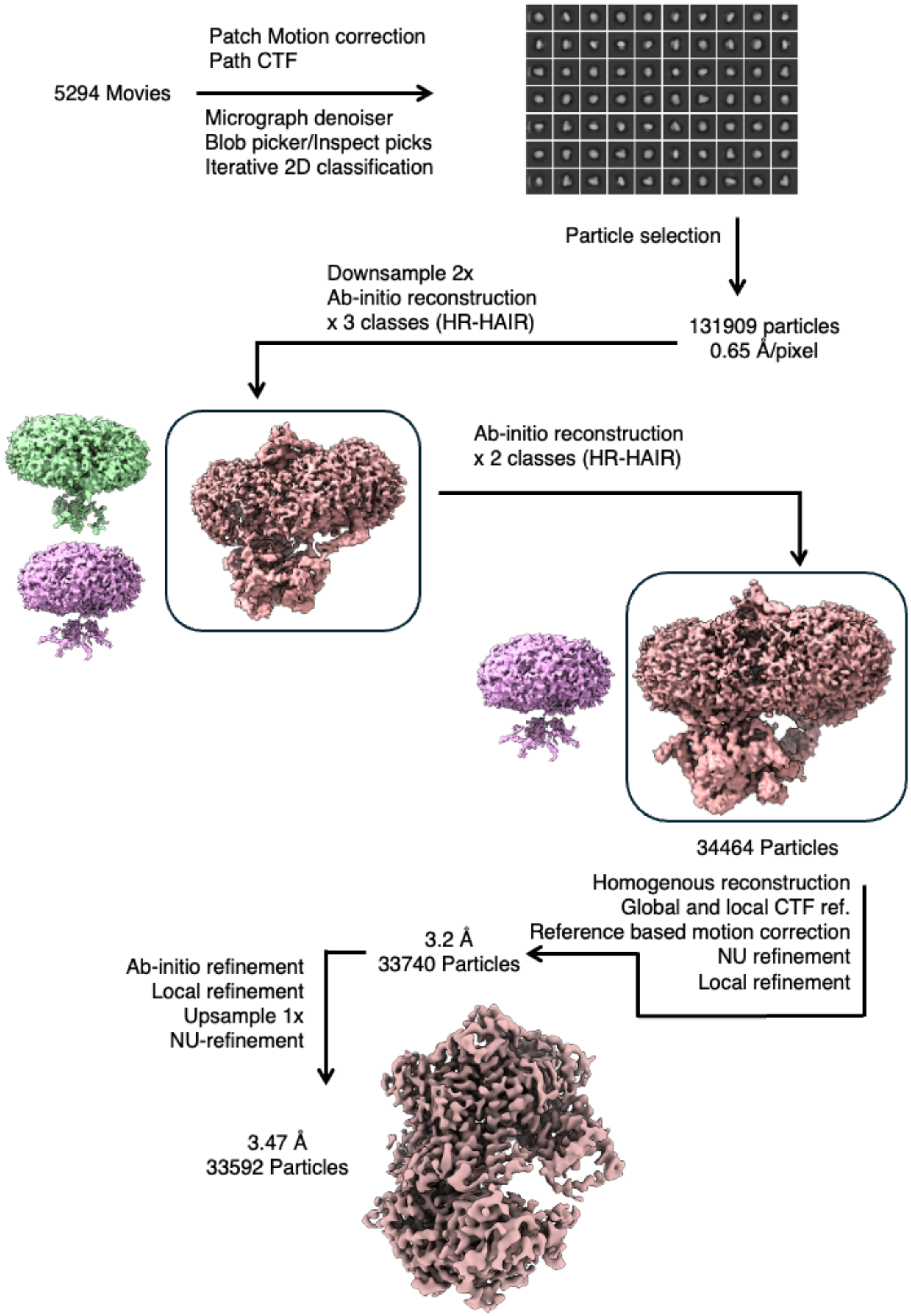

**Fig. S9. CryoEM workflow for post-catalytic NgFks1 apo**

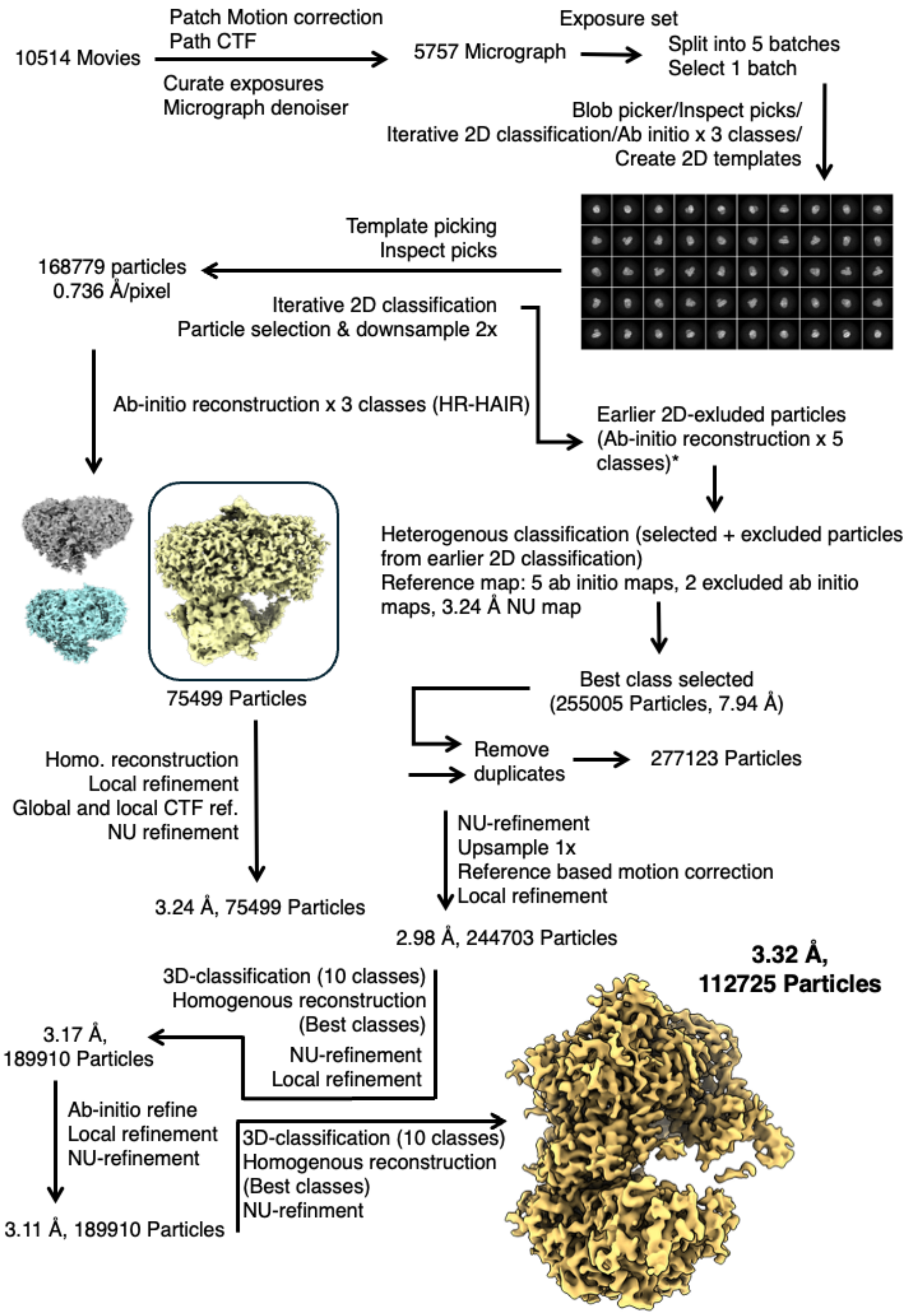

**Fig. S10. Cryo-EM workflow for the post-catalytic NgFks1 CFN**

| **Table S1.** Cryo-EM data collection and refinement statistics for **Post-catalytic NgFks1 apo** | |
|---|---|
| **Data Collection** | |
| Microscope | TFS Krios |
| Detector | Gatan K3 BioContinuum (6k x 4k) |
| Image pixel size | 0.65 Å |
| Defocus range | 0.8-2.2 µm |
| Electron exposure | 41 e-/Å² |
| Number of micrographs | 5294 |
| **Image Processing** | |
| Number of particles used | 33,592 |
| Symmetry | C1 |
| Final box size (pixels) | 512 |
| Reported resolution | 3.47 Å |
| FSC threshold | 0.143 |
| **Refinement** | |
| Atoms | 24,703 total; 12,315 non-H |
| Residues | 1,487 total; 1,465 protein |
| Water | 0 |
| Supplied resolution | 3.47 Å |
| B-factor (Å$^2$) | |
| Iso/Aniso (#) | 12315/0 |
| Protein (min/max/mean) | 25.01/170.73/80.11 |
| Length (Å) | 0.005 |
| Angles (°) | 0.900 |
| **Validation** | |
| MolProbity score | 0.91 |
| Clash score | 0.41 |
| Ramachandran plot (%) | |
| Outliners | 0 |
| Allowed | 3.90 |
| Favored | 96.10 |
| **Map-model fit** | |
| CC (mask) | 0.79 |
| CC (box) | 0.60 |
| CC (peaks) | 0.49 |
| CC (volume) | 0.77 |
| PDB ID | 11KD (Held for publication) |
| EMDB ID | EMD-75764 (Held for publication) |

| **Table S2.** Cryo-EM data collection and refinement statistics for Post-catalytic NgFks1 CFN | |
|---|---|
| **Data Collection** | |
| Microscope | TFS Krios |
| Detector | TFS Falcon 4i (4k x 4k) |
| Image pixel size | 0.734 Å |
| Defocus range | 0.6-2.4 μm |
| Electron exposure | 32 e-/Å² |
| Number of micrographs | 10514 |
| **Image Processing** | |
| Number of particles used | 112,725 |
| Symmetry | C1 |
| Final box size (pixels) | 512 |
| Reported resolution | 3.32 Å |
| FSC threshold | 0.143 |
| **Refinement** | |
| Atoms | 24,799 total; 12,309 non-H |
| Residues | 1,474 total; 1,443 protein |
| Water | 0 |
| Supplied resolution | 3.32 Å |
| B-factor ($Å^2$) | |
| Iso/Aniso (#) | 12309/0 |
| Protein (min/max/mean) | 4.37/127.79/49.02 |
| Length (Å) | 0.007 |
| Angles (°) | 0.970 |
| **Validation** | |
| MolProbity score | 0.94 |
| Clash score | 0.73 |
| Ramachandran plot (%) | |
| Outliners | 0 |
| Allowed | 3.33 |
| Favored | 96.67 |
| **Map-model fit** | |
| CC (mask) | 0.79 |
| CC (box) | 0.64 |
| CC (peaks) | 0.56 |
| CC (volume) | 0.53 |
| PDB ID | 35UT (Held for publication) |
| EMDB ID | EMD-77198 (Held for publication) |
